# Tissue-Adhesive Endoscopic Tattoo Inks for Precise Marking and Surveillance of Gastrointestinal Lesions

**DOI:** 10.64898/2025.12.01.691174

**Authors:** Mallikarjun Gosangi, Jordan R. Yaron, Subhadeep Dutta, Harsh Sant, Cole Wampler, Vanshika Singh, Maheshwaran Duraiyarasu, Trishita Chowdhury, Vikram D. Kodibagkar, Lee Chichester, Rahul Pannala, Kaushal Rege

## Abstract

Accurate marking of lesions and polyps is essential for effective management of gastrointestinal (GI) diseases, including colorectal cancer. Existing FDA-approved endoscopic carbon suspensions diffuse extensively within tissue and can extravasate intraperitoneally, compromising lesion localization, increasing fibrosis risk, and triggering inflammatory responses. These limitations reduce surgical precision and adversely affect patient outcomes. Here, we developed Tissue Adhesive Tattoos, generation 2 (TAT2) inks, composed of iron oxide nanoparticles encapsulated in bioadhesive polymers to improve spatial accuracy while maintaining high endoscopic contrast. TAT2 inks exhibit easy injectability, long-term stability, low macrophage inflammation, and strong spatial localization in ex vivo porcine intestinal tissue. Compared to clinical carbon ink, which spreads widely, TAT2 inks remained sharply localized with excellent contrast after submucosal injection in live mice and pigs. TAT2 also enabled visualization from both luminal and serosal surfaces and supported precise quadrant marking. Additionally, TAT2 inks were MR-compatible, expanding multimodality. Overall, TAT2 inks are biocompatible, non-inflammatory, precisely localized, and improve endoscopic marking for GI disease.

**Teaser:** Engineered bioadhesive tattoo inks deliver precise, durable GI marking with minimal spread and compatibility across imaging modalities.

## Introduction

In 2020, NIH allocated $3.1 billion (7.5% of its budget) for GI research. In 2018, expenditures for gastrointestinal healthcare reached $119.6 billion (*1*). In the same year, gastrointestinal endoscopies numbered 22.2 million, and 284,844 new cases of gastrointestinal cancers were diagnosed (*1*). Annually, nearly 11 million colonoscopies and more than 6 million upper endoscopies are performed in the United States (*2*). In 2023, the American Cancer Society estimated 153,020 colorectal cancer (CRC) diagnoses and 52,550 deaths in the United States. Approximately 19,550 diagnoses and 3,750 deaths will occur in individuals under 50 (*3*), highlighting the need for increased awareness and early detection.

Endoscopic tattooing is one of the most widely adopted techniques for marking gastrointestinal (GI) lesions to facilitate localization, subsequent surgical resection, and surveillance(*4*). Especially in colorectal cancers (CRC), it is crucial to identify potentially malignant lesions in the early stages of the disease, followed by resection, to improve patient survival rates (*5*). In addition, because surgeons often face challenges due to the absence of tactile sensation during laparoscopic procedures where primary lesions are difficult to visualize on the serosal surface of an organ such as the colon or bladder, mucosal markers that are visible on the serosal aspect are utilized to enable precise localization and achieve negative margins (*6, 7*) (*8*). Marking gastrointestinal lesions with tattoo ink is recommended by the United States multi-society task force (*9*) and is supported by significant clinical benefit. Polypectomy using colonoscopy reduces the incidence of and mortality by up to 60% compared to population control, as it removes early-stage adenomas (*10*). Endoscopic marking of advanced adenomas at the index colonoscopy facilitates lesion identification for subsequent endoscopic mucosal resection or submucosal dissection. Lesional marking is particularly recommended for subtle lesions such as sessile serrated adenomas or mucosal dysplasia in inflammatory bowel disease. The guidelines recommend marking 3-5 cm away on either side of the lesion but this can present clinical challenges in accurate identification. In Urology, identifying lesions with exact margins using a dye (e.g. India ink) or fiducial marker (e.g. TraceIT) implantation surrounding the tumor bed reduces the time and complexity of the transurethral resection of bladder tumors (TURBT)(*11*). Moreover, failure to precisely mark increases the risk of inadvertent excision of a healthy tissue or positive surgical margins, with elevated risk for cancer recurrence and the need for repeat procedures with increased risk of morbidity and mortality (*12, 13*).

Several materials have been utilizedfor preoperative localization in medical procedures; however, none satisfactorily addresses the diffusion problem. These materials include methylene blue(*14*), indigo carmine(*15*), toluidine blue(*16*), Indocyanine green (ICG)(*17*), and India ink(*18, 19*). Despite extensive research to identify a better contrast agent with improved digestive tract persistence, only one ink continues to be approved by the FDA: a purified carbon particle suspension (SPOT, GI Supply – now Laborie, Camp Hill, Pennsylvania)(*20, 21*). However, SPOT®, and its successors SPOT® Ex, etc., exhibit extensive tissue diffusion (*22*), clinical evidence of inflammation(*23*), and deep infiltration into tumor tissue, resulting in fibrosis (*15, 24*). In some studies, as much as 18% (9 out of 50 cases) of tattoos made with India ink were inadequately detectable, as previously reported (*13, 25*). Furthermore, performance limitations of such inks may have surgical consequences and in one study it was reported that 8% (4 out of 50 cases) of visible tattoos created with India ink to be inaccurate during surgical operations(*26*). In other studies, there have been documented 14% inaccuracies in localization, even when various endoscopic tattooing materials were utilized(*26, 27*). As such, there remains a persistent challenge in colonoscopy to improve both the precision and the visualization of endoscopic marking inks. Further, current clinical inks are not radio-opaque and are therefore not visualized on noninasive imaging modalities such as magnetic resonance imaging or computed tomography.

We hypothesized that these challenges stem from inherent limitations of nanosized suspensions of bare carbon particles, which lack sufficient tissue adhesive properties. Here, we sought to enhance the tissue adhesive capabilities of endoscopic tattooing materials. Developing such advanced endoscopic markers is crucial because no material-based approaches have been explored to tackle this substantial problem effectively. We examined various colored metal nanoparticles, including Prussian blue, CuO, and Iron oxide. The black color of iron oxide metal particles met the required intensity under white light. Then, we started using iron oxide particles as a core, and to achieve a stable enough colloidal suspension without any leaching problems in the submucosal colon, we hypothesized that double encapsulation using biocompatible polymers, such as dextran sulfate, followed by chitosan, would be necessary. These double-coated magnetic iron oxide particles also possess MRI potential, which will be an additional benefit of using these particles as contrast agents to visualize the site of interest under non-invasive procedures.

## Results

### Synthesis and Characterization of Polymers

Deacetylated chitin, also known as chitosan, is a derivative of the naturally occurring biopolymer called “chitin”, a major component of the exoskeletons of crustaceans (*28, 29*). Chitosan is a mucoadhesive polycationic biopolymer, which is soluble at pH ≤ 6.5 because of the protonation of primary amines available from the deacetylation of chitin (*30, 31*). To explore the different physicochemical properties of the chitosan component, we derivatized chitosan with glycidyl trimethylammonium chloride (GTMAC) (*32*) to generate biopolymers with low (q1C), moderate (q2C), and high (q3C) degrees of quaternization. Chitosan is a poly-(*β*-1-4)-D-glucosamine that contains primary amines, produced by the deacetylation of chitin, a polymer of *β*-1-4-*N*-acetyl-D-glucosamine. These freely available primary amines react with the epoxide ring of GTMAC, producing N-(2-hydroxypropyl)-3-trimethylammonium chitosan chloride. The derivatization of chitosan with GTMAC was verified using ^1^H NMR, FTIR, ninhydrin, and polymer weight analysis using size exclusion chromatography (SEC). Moreover, the degree of quaternization (DQ%) was estimated using conductometric titration with a standard silver nitrate (AgNO_3_) solution. Proton NMR spectroscopy, as represented in Figure S1a (supporting information), showed a peak at 2.3-2.5 ppm corresponding to the methyl protons of the N-acetyl moiety of glucosamine and a peak at 3.45-3.65 ppm corresponding to the trimethylammonium hydrogens. The latter peak increased amplitude from q1CH to q3CH and was absent in the parental polymer (HMC). The small peak observed at the same range in spectra of the parental chitosan (HMC) biopolymer can be attributed to the hydrogen of the second carbon in the glucosamine ring (*33*). The peak from 5.3 - 5.5 ppm corresponds to the hydrogen of the first carbon of the glucosamine ring (z) where it was attached to GTMAC. In addition, peaks at 5.1-5.3 ppm and 4.8-5.1 ppm were characteristic of the first carbon hydrogens of glucosamine with primary amine (y-z) and glucosamine with *N*-acetyl amine (x), respectively. Taken together, these analyses confirm the derivatization of the parental chitosan biopolymer with GTMAC and are in good agreement with previous reports (*33–35*). Additionally, the conjugation was assessed using FT-IR spectroscopy (Figure S1b, supporting information), which exhibited a characteristic peak at 1475 cm^-1^. This peak was absent in the parental polymers and can be attributed to the appearance of C-H bending of trimethylammonium moieties (*32, 36*) present on the GTMAC-conjugated chitosan. This peak’s intensity increased with the increasing extent of conjugation from q1C to q3C. Furthermore, the consumption of reactive primary amines, resulting from conjugation with GTMAC, was estimated using the Ninhydrin assay (*37*). The assay (Figure S1c, supporting information) confirmed the conjugation by showing the reduction of primary amines in the polymers in the order of HMC (100%) > q1C (∼75%) > q2C (∼60%) > q3C (∼30%). Moreover, the degree of quaternization (DQ%) was calculated by estimating the amount of chloride ions present on the conjugated GTMAC groups using conductometric titration of a chitosan solution against a standard AgNO3 solution (*38*). Figure S1d (supporting information) demonstrates that the percentage of degree of quaternization q3C (∼57%) > q2C (∼26%) > q1C (∼11%). Finally, the molecular weight determination of these polymers using gel permeation chromatography (GPC) indicated the molecular weight of HMC as 239 kDa < q1C as 253 kDa < q2C as 273 kDa < q3C as 278 kDa (Figure S2, supporting information), which further is indicative of the derivatization of HMC with the quaternary ammonium moieties.

### Synthesis and Characterization of TAT2 Inks

A key design requirement for effective GI inks is demonstrating high contrast under illumination with wide-spectrum, white endoscopic light and adhering to tissue. Given this requirement, the dark color of iron oxide (Fe_3_O_4_) nanoparticles makes them suitable for use as GI inks. Furthermore, iron oxide nanoparticles are considered safe and have been approved by the FDA for human use [38], and can be utilized for magnetic resonance (MR) imaging, thereby imparting multimodal imaging capabilities to these new inks. We employed dextran sulfate (DS; MW: 40kDa), which is a biocompatible synthetic sulfate polysaccharide for templating the formation and coating of iron oxide nanoparticles using a one-pot coprecipitation method (*39*) with slight modifications (**Figure 1**). Transmission electron microscopy (TEM) and dynamic light scattering (DLS) data of these anionic, black-colored DS-coated iron oxide nanoparticles (IO) dispersion indicated polymer (DS) coating surrounding several spherical core iron oxide nanoparticles ranging 8 – 20 nm in diameter and Z-average hydrodynamic diameter was 240 ± 10 nm (PI: 0.17 ± 0.028) and the zeta potential was-40 ± 5 mV at a pH ∼9.5, respectively (**Figures S3a and S3b, supporting Information**). The X-ray diffraction (XRD) spectra of IO exhibit characteristic signals at 220, 311, 400, 422, 333, and 440, corresponding to the crystallographic planes of Fe_3_O_4_ (**Figure S3c, Supporting Information**), which is consistent with previous reports.(*40, 41*) These results demonstrate that the dextran-coated iron oxide with Fe_3_O_4_ (magnetite) was successfully achieved.

**Figure 1.**
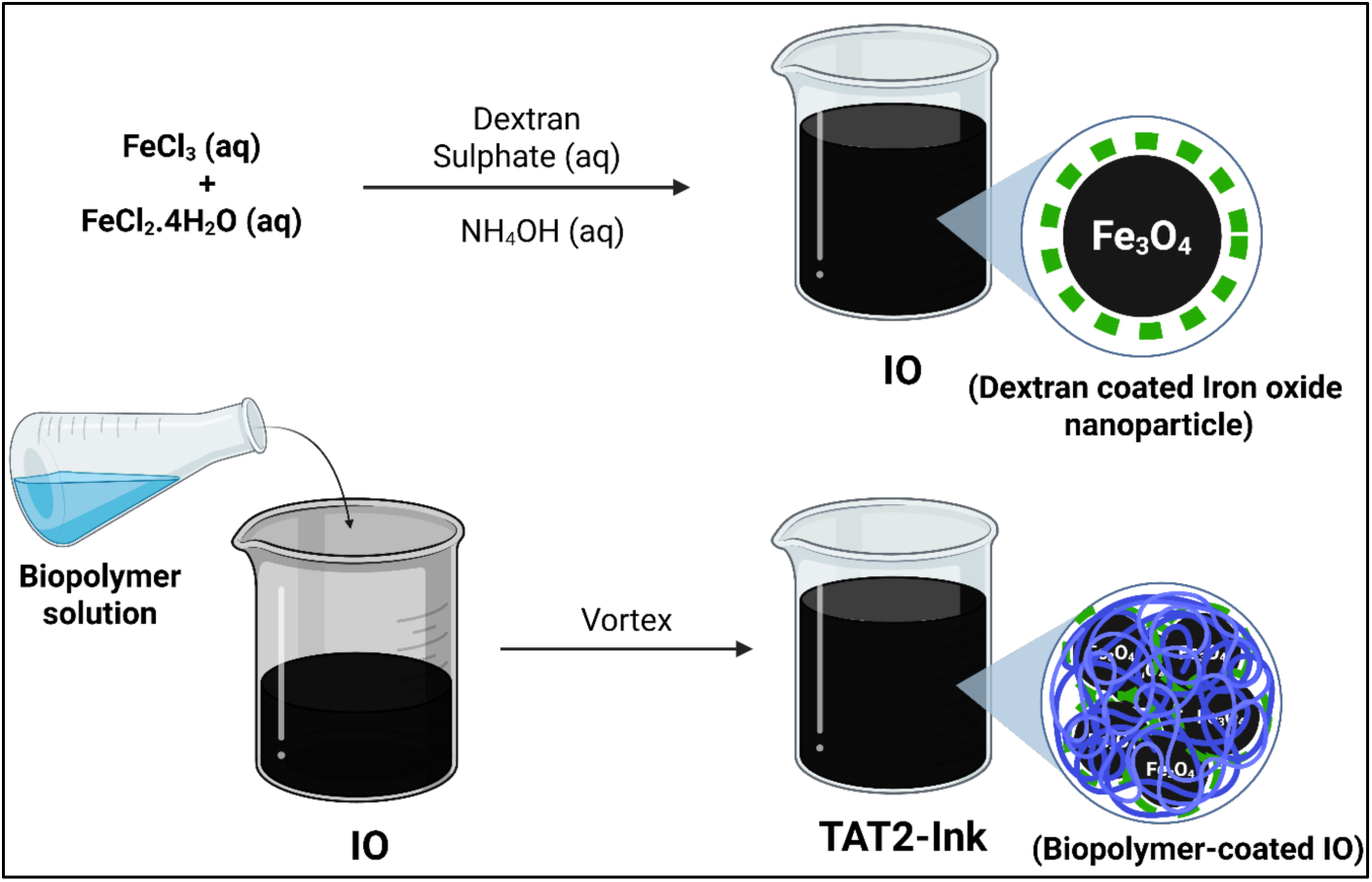
Schematic of the generation of dextran-coated iron oxide nanoparticles (IO) followed by coating with chitosan biopolymer, resulting in TAT2 inks.

Cationic, tissue-adhesive biopolymers such as HMC or quaternized chitosans were complexed with the anionic IO at different ratios (0.5:1, 1:1, 2:1, and 3:1) to generate different formulations of TAT2 inks. The Z-average hydrodynamic diameters of these TAT2 inks remained invariant beyond a 1:1 ratio (**Figure S4, Supporting Information**); subsequent TAT2 ink formulations were generated using this ratio. Z-average hydrodynamic diameters and zeta potential values were 1070 ± 256 nm (PI: 0.5 ± 0.03) and +32 ± 12 mV for hC-IO, 1267 ± 59 nm (PI: 0.12 ± 0.06) and +39 ± 8 mV for q1C-IO, and 1674 ± 467 nm (PI: 0.15 ± 0.1) and 52 ± 6 mV for q3C-IO, respectively (**Figure 2a and 2b**). The Z-average hydrodynamic diameters of TAT2 inks were substantially larger than those of the clinical ink and IO particles, likely due to the encapsulation of multiple IO particles by the cationic biopolymers. Furthermore, the change in the negative zeta potential of anionic IO particles to positive values for cationic TAT2 inks is attributed to the larger dimensions and cationic nature of these inks, which we reason will facilitate higher retention in the tissue upon endoscopic administration (*42*).

**Figure 2.**
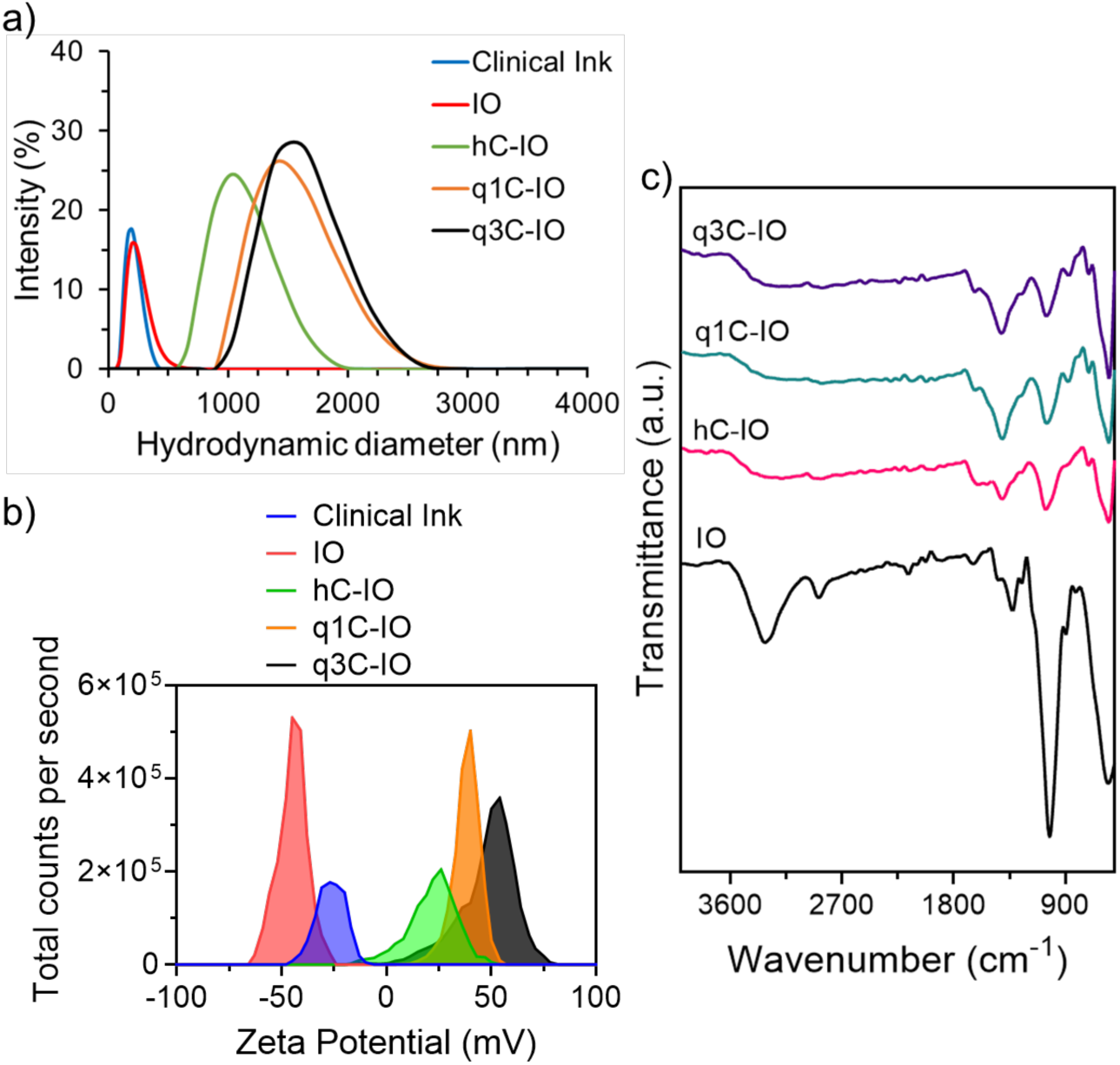
Characterization of TAT2 inks. a) Hydrodynamic diameters (nm), (b) Zeta potentials (mV) of dextran sulfate-coated iron oxide nanoparticles (IO: red), hC-IO (green), q1C-IO (orange), and q3C-IO (black) in comparison to the clinical ink (blue). c) Representative FTIR spectra of n = 3 independent ink formulations.

FTIR spectra of IO (**Figure 2c**) exhibit strong peaks at 3325 cm^-1^ and 2910 cm^-1^, indicating the presence of OH, NH_2,_ and CH– vibrations. These confirm the moieties present in dextran sulfate-coated iron oxide nanoparticles. In the case of cationic biopolymer-coated IO nanoparticles, peaks at 1634 cm^−1^ (N-H vibration) and 1405 cm^−1^ (C-N vibration) indicated the presence of chitosan or its quaternized derivatives. The peak at 1024 cm^−1^, indicative of ether moieties, was seen in all inks, as is expected from the polymer coating in these particles (*43–45*). Finally, all spectra exhibit a peak at 560 cm^-1^, indicating Fe-O stretching in iron oxide (*41*). Scanning Electron microscopy (SEM) images of both IO and q3C-IO particles in **Figure 2d** demonstrate that both particles are spherical in morphology, showing dextran-coated iron oxide (IO) nanoparticles becoming larger after complexing with the chitosan polymer coating (q3C-IO). The particles of q3C-IO that appeared in the SEM image were further confirmed with the Energy-dispersive X-ray (EDX) spectroscopy using a line scan method, and elemental analysis to identify them as polymer-coated iron oxide particles (**Figure S5, Supporting Information**). Elemental analyses using line scan mode indicated that the spherical particle has an enriched iron oxide core surrounded by a carbon layer. (*45–47*).

### Shelf-life stability and Injectability

The proximal and distal colon exhibit pH ranges from 5.5 to 6.5 and 6.5 to 7.0, respectively.(*48*) Chitosan-IO (hC-IO) inks demonstrated excellent colloidal stability at pH 6.5 but not at pH 7.4 (Figure S6a, Supporting Information). However, quaternized chitosan-IO (q3C-IO) TAT2 inks were stable at pH levels of 6.5, 7.0, and 7.4 (Figure S6b, Supporting Information), likely due to the permanent cationic charge from the trimethylammonium groups on the polymer, which ensures stability under various conditions. Considering the environment in the colon, inks at pH 6.5 were used in all subsequent studies. TAT2 ink (q3C-IO) demonstrated excellent shelf stability over at least three months when stored at 4°C in a degassed desiccator; no loss in hydrodynamic diameter (Figure 3a) or zeta potential value was observed during this period due to the strong cationic charge from q3C polymer whereas other TAT2 inks (hC-IO, and q1C-IO) lost their hydrodynamic diameter after a month or two. Colon tattoos are delivered through a long endoscopic needle (2.4 m) into the submucosa of the intestine, which requires that ink formulations be stable and easily injectable.(*49, 50*) We evaluated the effect of shear and gauge of the endoscopic needle on the colloidal behavior of TAT2 inks, which was maintained (Z-average hydrodynamic diameter: Figure 3b and zeta potential: Figure 3c) before and after passing through the same endoscopic needle used for the endoscopic injection in the clinic and exhibit viscosities similar to those of water and Clinical Ink at higher shear rates, and their viscosities remain invariant with shear, indicating Newtonian behaviour (Figures 3d). Even hydrogels that possess higher viscosities than water have been injected into the intestinal submucosa (*49, 50*) and TAT2 inks with viscosities similar to water are easily injectable.

**Figure 3.**
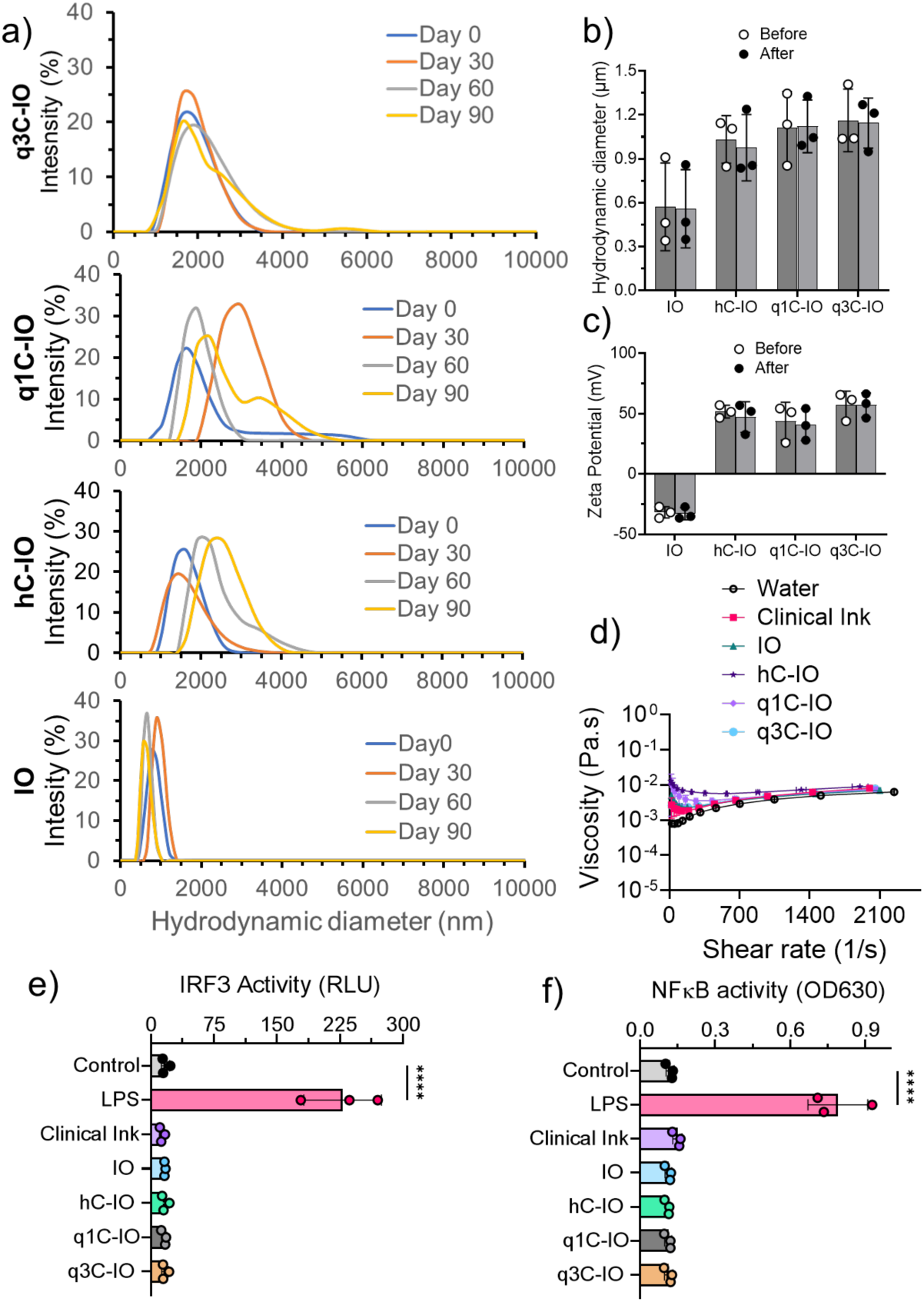
Shelf-life stability and injectability of TAT2 inks. **a)** hydrodynamic diameter of TAT2 inks on the day of preparation (blue line), one month (orange line), two months (grey line), and three months (yellow line) stored at 4 °C under vacuum. **b).** Hydrodynamic diameter (μm) and **c)** Zeta potential (mV) of IO, hC-IO, q1C-IO, and q3C-IO TAT2 inks before and after injecting via an (Interject^TM^, CONTRAST, Boston Scientific, 23 gauge, and 2.4m) endoscopic loop and needle (n=3). **d)** Viscosity (Pa.s) as a function of shear rate (1/s) of different TAT2 inks in comparison with water and clinical ink (n=3) at room temperature (∼25 °C). **(e)** Activation of IRF3, measured by secreted luciferase, and **(f)** NF-kB, measured by secreted alkaline phosphatase for individual TAT2 inks and clinical ink after 24 hours of incubation. Lipopolysaccharide (LPS) treatment and unstimulated cells were used as positive and negative controls, respectively. Statistics were calculated as one-way ANOVA with Fischer’s LSD using untreated cells as the comparison group. In the figure, ****p<0.0001.

### Evaluation of the Inflammatory Activity of TAT2 Inks *in vitro*

Tissue-resident macrophages are the earliest cell population to participate in the foreign body response. Biomaterial design strategies are widely recommended to take macrophage responses as key considerations for biocompatibility.(*51*) We utilized the J774-DUAL^TM^ murine macrophage cell line, a reporter cell line for IRF3 and NF-ᴋB transcription factor activation, to determine the inflammatory potential of our TAT2 formulations (*52, 53*). Parental (HMC) and quaternized (q1CH, q2CH, and q3CH) chitosan did not activate IRF3 and NF-ᴋB signalling (**Figure S7**, **supporting information section**). Similarly, Clinical Ink, hC-IO, q1C-IO, and q3C-IO TAT2 inks demonstrated negligible inflammatory activity in the J774-DUAL^TM^ cell assay (**Figure 3e and 3f**). These results suggest that the formulated TAT2 inks have minimal potential for acute inflammatory stimulation.

### Investigation of TAT2 inks ex vivo

There are two submucosal injection methods used in the clinic to successfully place tattoos while avoiding peritoneal spill or unintended deposition in extra-colonic tissues such as the omentum, small intestine, or kidney(*54*). The first method involves creating a saline bleb by injecting 0.5–1 mL of saline into the submucosa, followed by the insertion of the tattoo needle into the bleb to deliver the ink. The second method, commonly used by expert endoscopists, involves approaching the submucosa tangentially (rather than *en face*), inserting the needle into the submucosa, lifting it toward the center of the lumen, and delivering the ink. Following the second method used clinically, we adopted an injection technique to evaluate our inks ex vivo using fresh porcine tissues (**Figure 4a)**. The clinical ink or TAT2 inks were injected into 5 cm x 3 cm sections of porcine intestines ex vivo (**Figure 4b**). The area and contrast (mean grey value) of the ink in the tissue were determined 30 minutes post-injection, as described in the Experimental section. The contrast of all TAT2 inks was similar to that of the clinical ink (**Figure 4c** and **4d**), indicating similar performance for visualization under white light. However, the area of the TAT2 inks was significantly lower compared to that of the clinical ink (**Figure 4d**), indicating greater fidelity of spatial localization with the new inks. Increasing the concentration of TAT2 inks from 1.25 to 10 mg/mL did not result in significant changes in contrast to tissue under white light, but led to a somewhat higher area. However, this was still significantly lower than the clinical ink (**Figure S8, supporting information section**). We continued to use 10mg/mL for inks to ensure sufficient contrast in eventual in vivo applications. Investigation of diffusion in tissue over time indicated that TAT2 inks did not change localization significantly; however, the clinical ink continued to diffuse and spread over the 24 hours evaluated. The area of the clinical ink almost doubled from ∼0.7 cm^2^ immediately upon injection to ∼1.4 cm^2^ at 24 h. At all times, the contrast of the individual TAT2 inks was similar to that of the clinical ink, indicating no loss in visualization. These results suggest a significant benefit in tissue marking with TAT2 inks and underscore the significant challenges seen with the only FDA-approved clinical ink.

**Figure 4.**
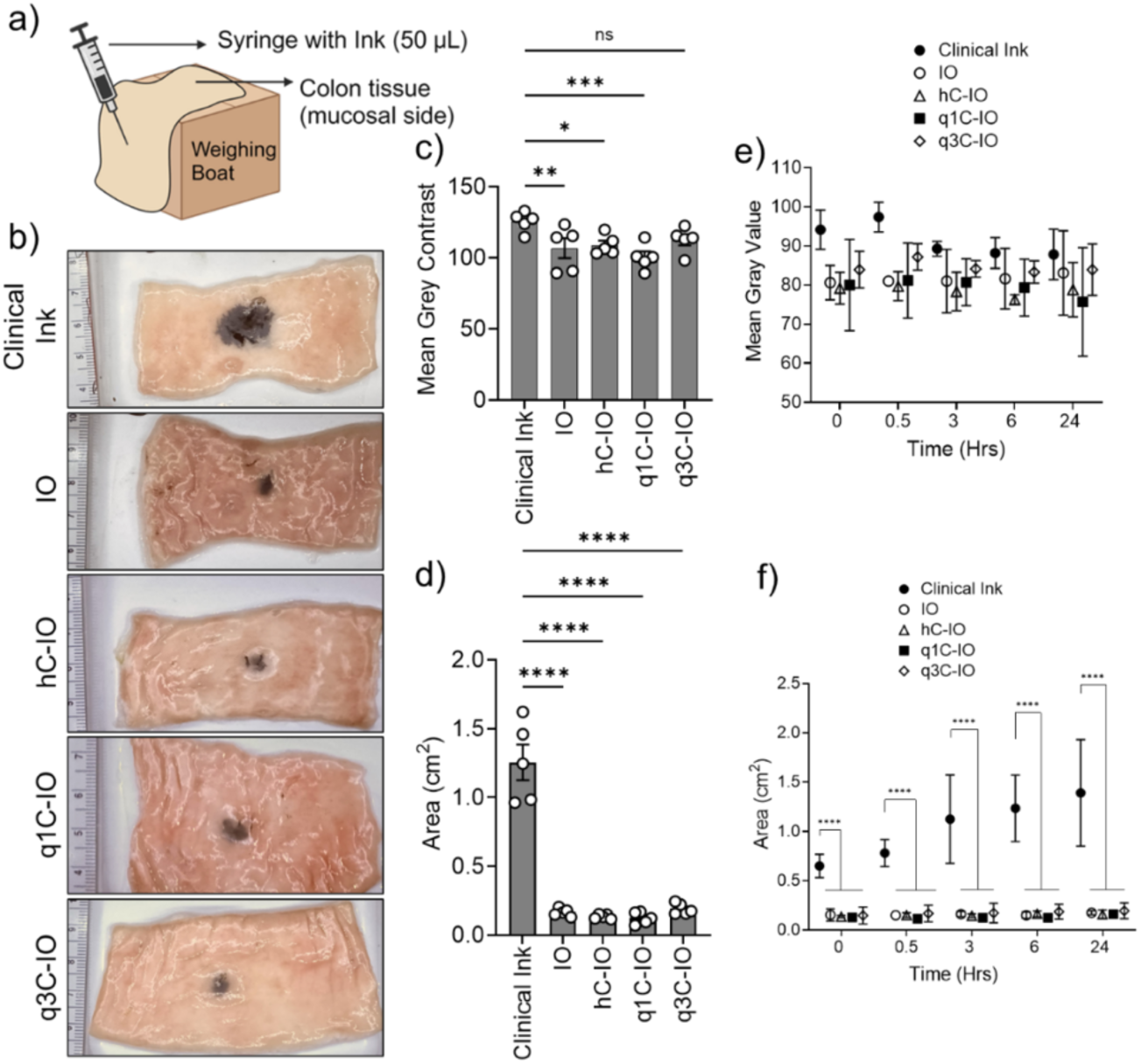
Evaluation of TAT2 ink performance in porcine colon tissue ex vivo: **a)** Schematic representation of injection (∼10° incident with respect to tissue surface) in porcine colon tissue ex vivo to mimic the endoscopic injection in clinical practice. **b)** Representative images of colon tissue injected with 50 mL of IO, hC-IO, q1C-IO & q3C-IO TAT2 inks or the clinical ink. **(c)** Quantified mean grey contrast value and **(d)** Area (cm^2^) of TAT2 inks (1:1 chitosan:IO, 10mg/mL) or clinical ink (n=5). Characterization of diffusion of TAT2 inks or clinical ink in porcine tissue over time as determined using **(e)** contrast in tissue to white light and **(f)** Area (cm^2^) (n=3). Statistical significance was calculated using one-way ANOVA with Fischer’s LSD using the clinical ink as group for comparison. In the figure, *p<0.05, **p<0.01, ***p<0.001, ****p<0.0001.

### Evaluation of Efficacy and Biocompatibility of TAT2 inks using Subcutaneous Administration in Live Mice

Sub-cutaneous implantation in immunocompetent mice is commonly used for determination of biocompatibility of nanomaterials and is consistent with FDA recommendations for evaluation of medical devices (*55–57*). Four TAT2 inks - IO, hC-IO, q1C-IO, q3C-IO – and the clinical ink were injected into subcutaneous region of in the dorsum of immunocompetent BALB/c mice as shown in **Figure. 5a**. Two injections, at least 1 cm apart, were made with two separate inks (50 µL per injection) for which the endotoxin content was verified to be < 0.5 EU (endotoxin units)/mL within 24 hours of in vivo studies (*58*). TAT2 inks were administered at a dose of 10 mg/mL based on ex vivo optimization studies. The equivalent iron concentration for this dose was 6-7 mg/mL of iron (ICP-MS analyses), which in turn translated to 20 mg/kg of iron per mouse - significantly below the toxicity threshold of 200 mg/kg of Fe for iron oxide nanoparticles (*59*). No mice in the study showed signs of distress, behavioural changes, inflammatory response, or adverse reaction at the injection site. The clinical ink exhibited extensive diffusion within the skin almost immediately and at 28 days the entire region appeared to be saturated with the ink (**Figure 5a**). By comparison, TAT2 formulations exhibited superior spot localization with minimal diffusion at 28 days post-implantation.

**Figure 5.**
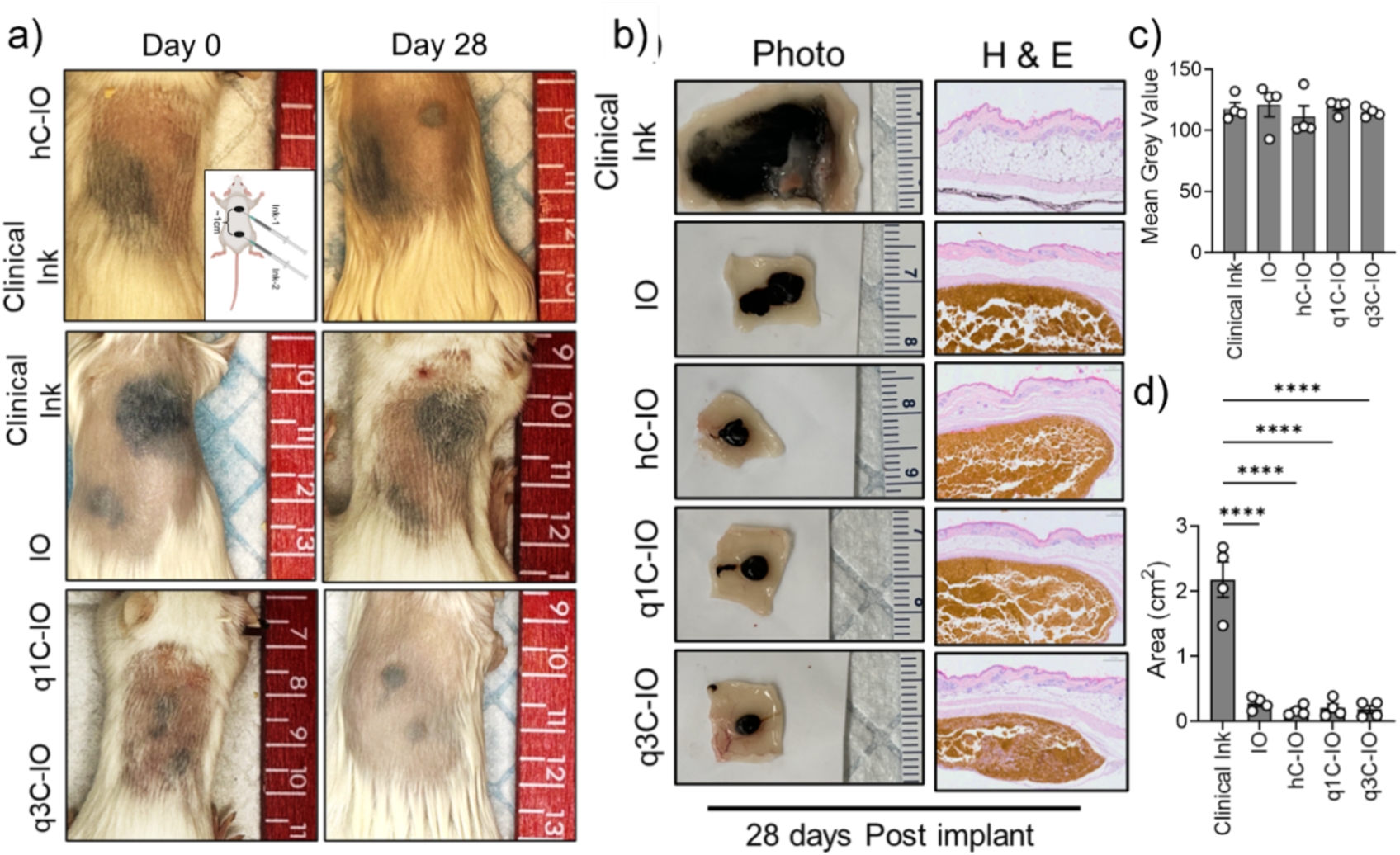
Efficacy of TAT2 inks in immunocompetent Balb/c mice. a). Representative images from immunocompetent Balb/c mice subcutaneously implanted with IO, hC-IO, q1C-IO, and q3C-IO TAT2 inks and the clinical ink in the sub-cutaneous space on day 0 (immediately after injection) and day 28. N=4/group. **b).** Representative images of skin tissue subcutis illustrating localization and area spread of inks in the subcutaneous space. **c).** Mean grey value of contrast and **d).** area covered by ink in skin tissue collected on day 28. Statistics were calculated as one-way ANOVA with Fisher’s LSD using the clinical ink as the group for comparison. *p<0.05, **p<0.01, ***p<0.001, ****p<0.0001.

Cellular responses and tissue retention of inks were assessed by histological evaluation and quantifying the diffusion area of the ink in excised dorsal skin samples collected 28 days post-injection, as depicted in **Figure 5b**. Histological evaluation using haematoxylin and eosin (H&E) staining confirmed the presence of iron oxide-based tattoo particles, visualized in subcutaneous region as golden-brown deposits which is consistent with the previous report (*60*), demonstrating tattoo tissue retention. TAT2 inks displayed strong association within cells and maintained strong localization at the injection site with minimal diffusion in the skin (**Figure 5b**). Quantitative analysis of excised skin sample images revealed consistent contrast and diffusion patterns (**Figure 5c** and **5d**), replicating the observations made in the ex vivo study.

### Evaluation of TAT2 inks Efficacy in Live Pigs

Five inks - IO, hC-IO, q1C-IO, q3C-IO, and the clinical ink were injected into the sub-mucosa of live Yorkshire pigs using a therapeutic gastroscope and methods commonly used in humans (**Figure 6a**). Representative endoscopic images (**Figure 6b**) of the pig colon on day 0 (immediately after injection) and day 14 revealed that all TAT2 inks were easily detectable and maintained their localization with no local inflammatory response (**Movie S1, Pig Colonoscopy, Supporting information**). However, the clinical ink diffused extensively in the tissue and lost any spatial localization information, a feature that is integral to the function of endoscopic inks. In one of the four pigs, the clinical ink leaked into the peritoneum and diffused to the pancreas after injection into the colon, an event never observed with any of the TAT2 formulations. The q3C-IO TAT2 ink also demonstrated excellent localization on day 0 and day 14 when administered as a quadrant pattern of small spots in the colon (**Figure 6c**), a standard clinical procedure for marking lesions that often suffer from poor visualization due to the diffusion of existing clinical inks.(*54, 61, 62*) During necropsy at the 14-day follow-up, the spots were easily identified through an open abdominal incision (laparotomy). In the present study, the visibility (contrast) and precision (low diffusion) of TAT2 inks were assessed on the mucosal and serosal sides of the colon, followed by euthanasia of each animal. TAT2 inks demonstrated excellent visualization on both mucosal (luminal) and serosal sides of the colon (**Figure 6d**), and no spillage into the peritoneal cavity was observed. Even at 14 days, q3C-IO TAT2 ink maintained its localization in the quadrant, and no overlap was seen between individual spots (**Figure 6e**), on both serosal and mucosal sides of the intestine, which was further confirmed with the histological analysis (**Figure S9, Supporting information**) of both H & E and pression blue. TAT2 ink showed clear enhanced spot localization behavior compared to the clinical ink, which diffused over a large tissue region (**Figure 6b)**. These results, with the clinical ink in the submucosa of live pigs, are similar to those seen in live mice (**Figure 5a**) and reported in the literature (*63, 64*). The ability to detect inks on the serosal side is critical in laparoscopic surgeries, where tactile detection of polyps is not possible(*65, 66*). Further, inadvertent intraperitoneal spillage of endoscopically applied tattoo inks has been documented in up to 13% of reported cases (*67, 68*). This occurrence leads to a notable decrease in the visibility and identification of anatomical structures and planes during surgery. Quantitative analyses indicated that TAT2 inks demonstrated similar tissue contrast under white (endoscopic) light on mucosal and serosal surfaces (**Figure 6f and 6g**). All TAT2 inks appeared as a precise spot with a diameter of ∼1 cm. In addition, the area covered by the clinical ink upon injection (∼18 cm^2^) was approximately 70-fold higher than that covered by any TAT2 ink (e.g., q3C-IO: ∼0.25 cm^2^) on either side of the colon (**Figure 6f and 6g**), indicating inferior performance of the clinical ink in pigs using the same endoscopy equipment and methods used for humans.

**Figure 6.**
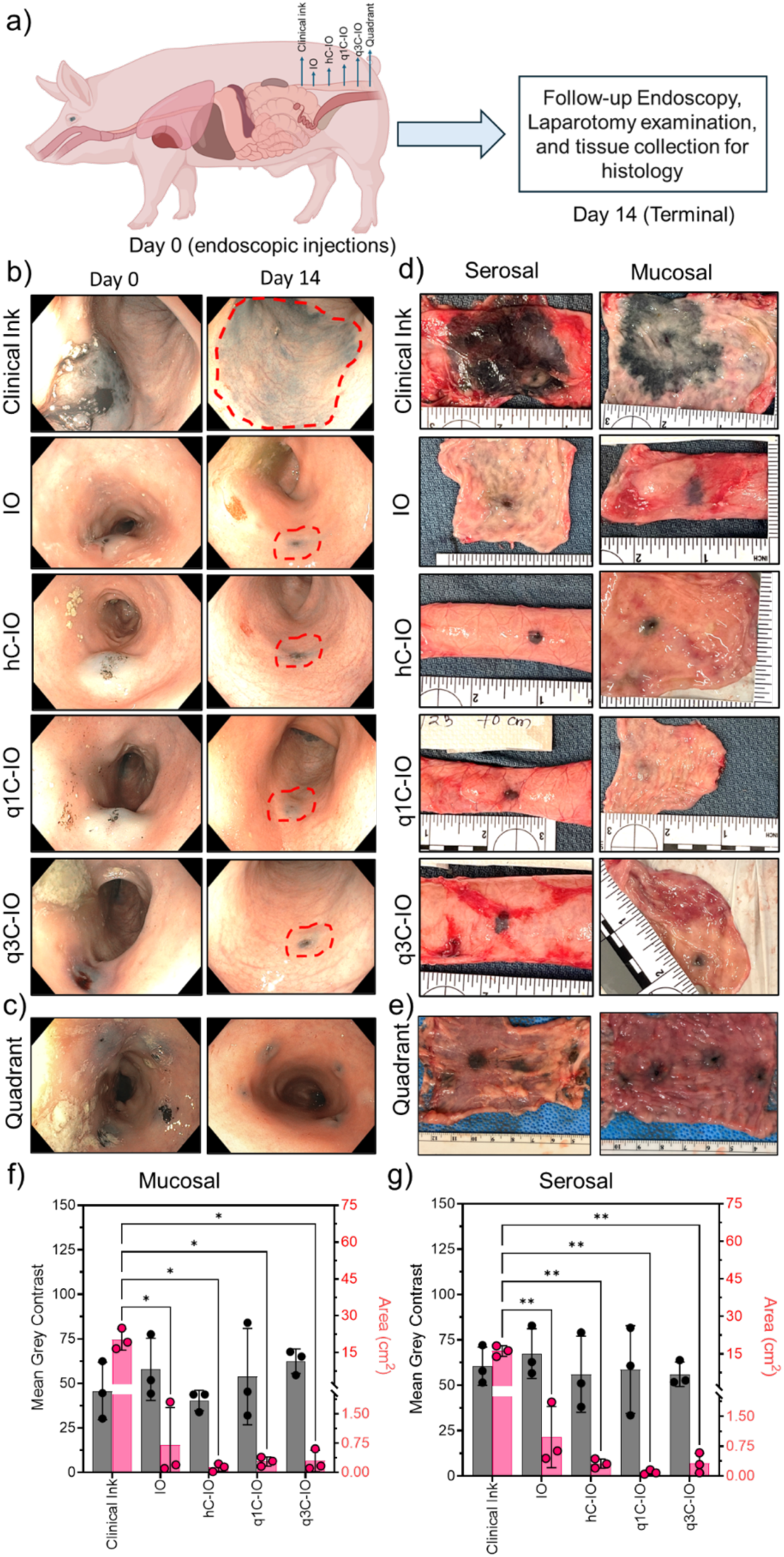
a) Schematic of the *in vivo* study plan of endoscopy evaluation of TAT2 inks in pigs during 2 weeks residence. **b)** Representative endoscopic images from n=5 of porcine colon on day 0 and day 14 after submucosal injection of 1 mL IO, hC-IO, q1C-IO, and q3C-IO TAT2 and the clinical inks, injected 5 cm apart. **c)** Endoscopic images on day 0 and on follow-up on day 14 of quadrantally placed q3C-IO TAT2 ink (0.5 mL per injection) to mimic clinical procedures used to mark lesions in the descending colon (n=3). **d)** Representative images of porcine colon tissue specimens showing TAT2 inks and clinical inks, and **e)** q3C-IO TAT2 ink quadrant on the mucosal and serosal side of the colon tissue collected on day 14 (n=3). Quantification of spot area (cm^2^), and contrast to tissue under white light (mean grey value) from **f)** mucosal (black) and **g)** serosal (pink) side in comparison to clinical ink (n=2). Statistics were calculated as one-way ANOVA with Fischer’s LSD using the clinical ink as a comparison group. In the figure, *p<0.05, **p<0.01, ***p<0.001, ****p<0.0001.

TAT2 q3C-IO was stable and uniform at physiological pH = 7.4 (**Figure S6, supporting information**) and used for further evaluation on extended residence time over 65 days in the Pigs’ colon. A total of four different sites in the colon (**Figure 7b**), separated by a 5 cm distance, were injected with one mL of freshly prepared q3C-IO, and a quadrant marking was circumferentially applied with 0.5 mL each at the most distal part of the colon. Videos (**Movie S2, supporting information**) and images (**Figure 7b**) were recorded on day 0, day 40, and day 65, indicating easy identification of all the locations injected with TAT2 and the persistence in precision in staying with no diffusion to merge each other at four 1 ml sites, and with quadrant marking. Clinical ink, when injected in a similar pattern, diffused, merged, and faded out, making it difficult to identify the locations (**Figure 7a**), including the quadrant. The tissues collected on day 65 (**Figure 8a**) were quantified for contrast (**Figure 8b**) and spot area (**Figure 8c**), and the results were compared with Clinical Ink. It indicates that the TAT2 after 65 days residence in the colon is equivalent, in contrast, and that the spot area is 99% less than the clinical ink. All four spots in the quadrant were dark and appeared with clear edges without any merging with each other. In contrast, the four spots of clinical ink in the quadrant merged into a single marking, indicating that a clear marking with multiple locations using the current clinical ink is impossible, unlike the TAT2.

**Figure 7.**
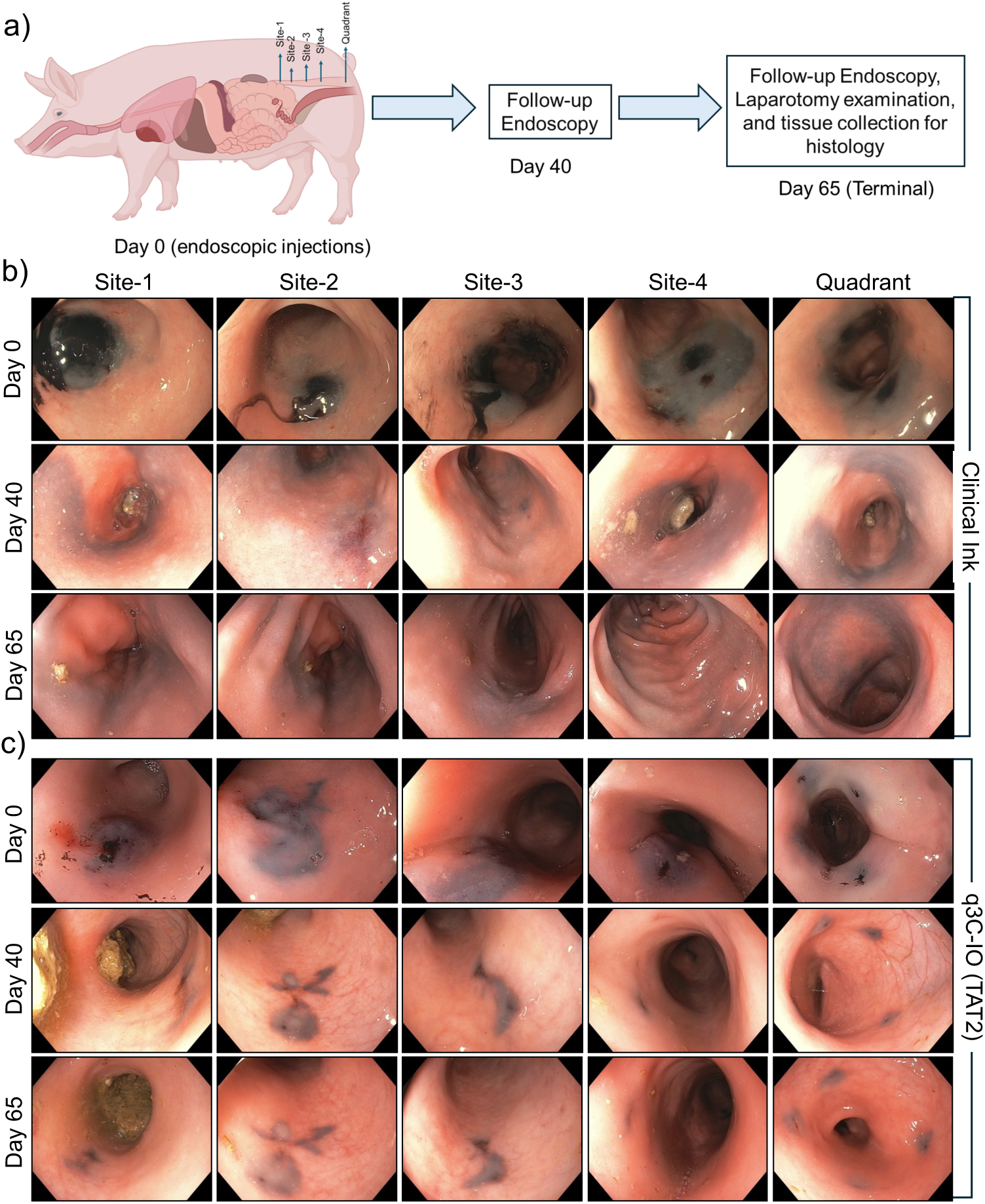
a) Schematic of the *in vivo* study plan of endoscopy evaluation of TAT2 inks in pigs during 65-days residence. **b, c)** Representative endoscopic images from n=5 of porcine colon on day 0, day 40, and day 65 after submucosal injection of q3C-IO TAT2 or the clinical ink, injected 5 cm apart. Sites 1, 2, 3, and 4 are single injection of 1 mL, while Quadrant indicates 4 injections of 0.5 mL spaced circumferentially around the colon.

**Figure 8.**
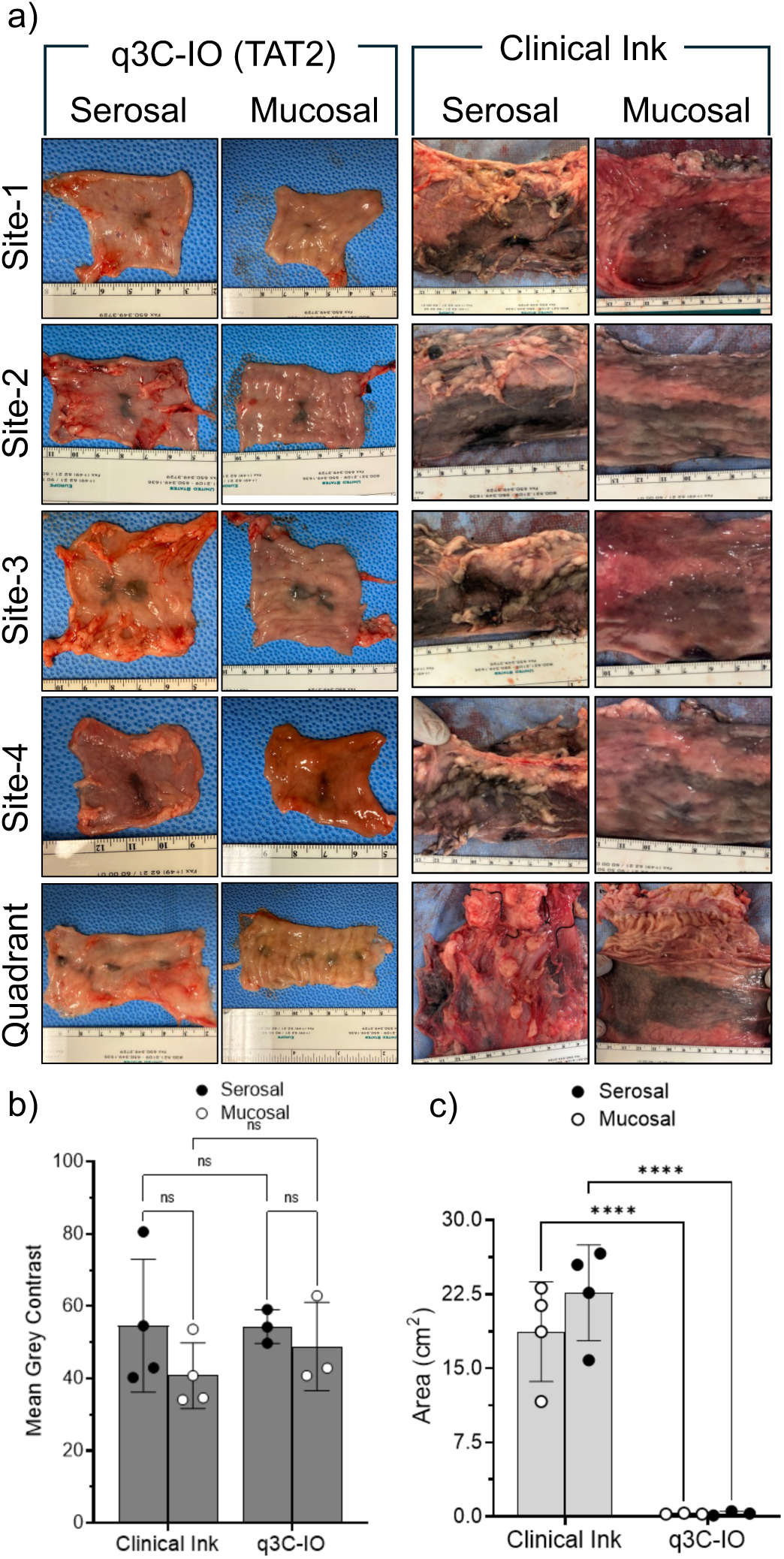
a) Representative images of porcine colon tissue specimens showing TAT2 ink q3C-IO and clinical ink for sites 1-4 and quadrant on the mucosal and serosal side of the colon tissue collected on day 65 (n=3-4). **b)** Quantification contrast to tissue under white light (mean grey value) and of **c)** spot area (cm^2^), from serosal (black) and mucosal (white open) side in comparison to clinical ink (n=3-4). Statistics were calculated as one-way ANOVA with Fischer’s LSD using the clinical ink as a comparison group. In the figure, *p<0.05, **p<0.01, ***p<0.001, ****p<0.0001.

### Histology and Evaluation of toxicity

Histological analysis with hematoxylin and eosin (H&E) staining revealed iron oxide-based tattoo particles as golden-brown deposits in the submucosal region [58], confirming their retention within the tissue. Further, Prussian blue staining is a histochemical technique for detecting and visualizing ferric iron (Fe3+), which we used to stain the same ion present in TAT2 as Fe_3_O_4_ (**Figure 9a, 10a**) (*69*). Evaluation of Prussian blue staining of TAT2 reinforces appreciation of the tight residence of TAT2 in the submucosal space of the colon without leaching to any adjacent locations, even after 14-day (**Fig 9a**) and 65-day (**Fig 10a**) span. To determine biocompatibility and toxicity after 14-and 65-days of residence in pigs, we performed histopathology of the end organs (liver, kidney, lung, and spleen) and quantified relevant clinical chemistries. Blinded reading of end-organ histology by a veterinarian with expertise in pathology determined no evidence of toxic or inflammatory tissue reactions. Lungs did not show extensive or diffuse inflammation or fibrosis, though one pig displayed minor focal sites of lymphoid activity, likely attributable to a pre-existing underlying exposure. Portal triads in the liver showed no inflammatory response, although minimal-to-mild congestion was observed in two animals, possibly occurring during postmortem tissue collection. Kidney cortices exhibited no adverse reactions, with no evidence of glomerulitis, capillaritis, or tubulitis. (**Figure 9b, 10b**). Importantly, no evidence of Prussian blue staining was observed in end-organs at 65-days, indicating that there was no detectable systemic migration of implanted material (**Fig 10b**). Spleen tissues were also evaluated, but due to the use of an injectable euthanasia agent, leading to an overwhelming presence of erythrocytes, it was deemed incompatible with full evaluation (not shown). Clinical chemistry evaluation of blood serum collected on day 0 (before implantation) and at 14-days post-implantation further supported biocompatibility with no elevations in serum creatinine (CRE; **Figure 9c**), blood urea nitrogen (BUN; **Figure 9d**), serum calcium (Sr. Ca^2+^; **Figure 9e**), or alanine transaminase (ALT; **Figure 9f**). Clinical chemistries for long-term implantation at day 0, 40, and 65 are provided in **Table S1** and demonstrate no significant or adverse responses of CRE, BUN, Sr. Ca^2+^, or ALT levels.. Complete blood count measures were also unaffected (not shown). Taken together, these findings indicate that in this study, neither TAT2 inks nor clinical ink (as these were simultaneously present in each pig) induced acute or short-term adverse pulmonary, hepatic, or renal toxicities.

**Figure 9.**
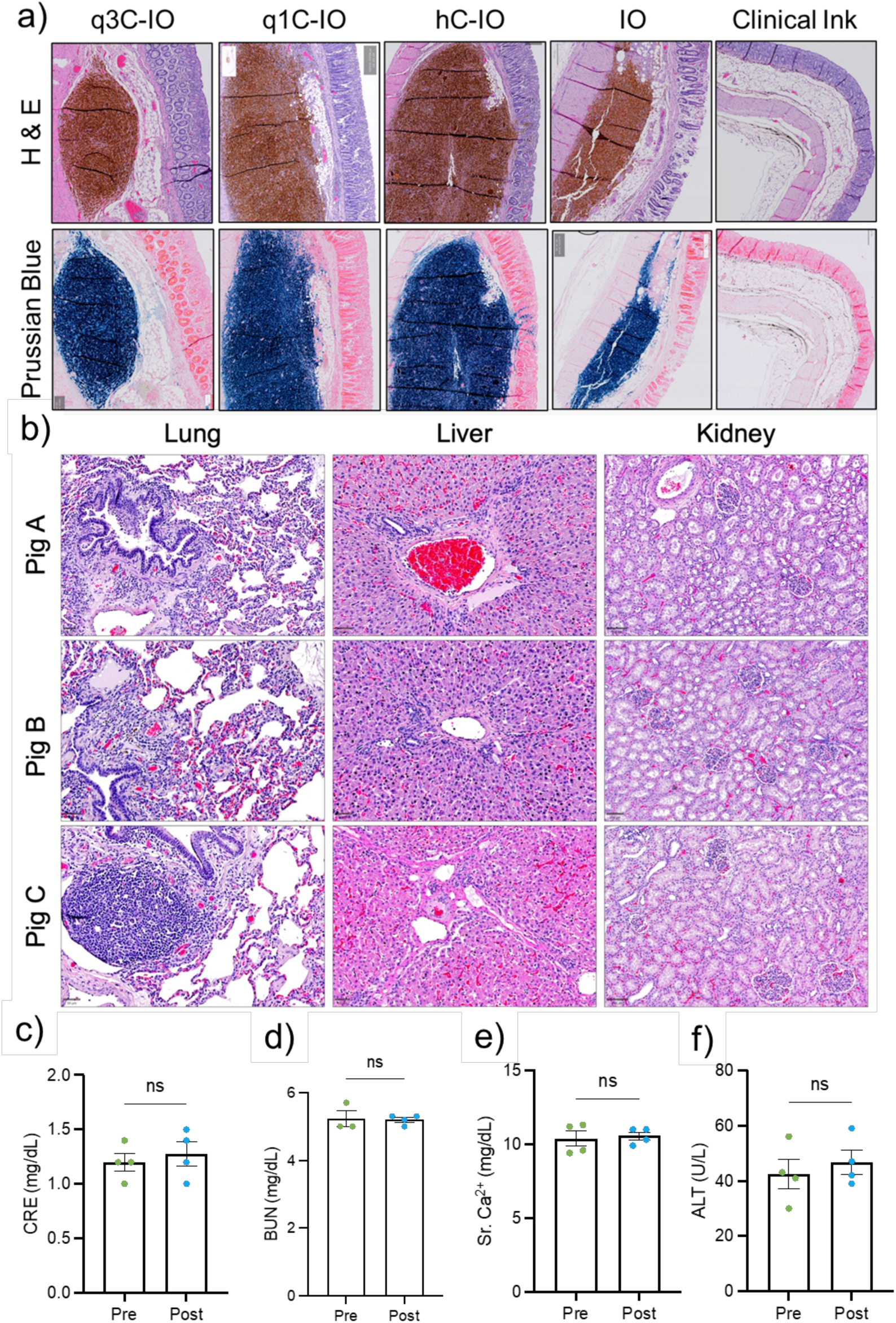
a) Representative histology images of H&E and Prussian blue staining of colon tissue administered with IO, hC-IO, q1C-IO & q3C-IO TAT2 ink and clinical inks after 14 days. (n=3) **b)** Representative H&E histology of pig end-organs 2 weeks post-implantation of clinical ink and TAT2 inks. No evidence of fibrosis or inflammation in the lung, inflammation or necrosis at the hepatic triad, or glomerulitis, capillaritis, or tubulitis in the kidney cortex was observed by blinded reading. Scale bar is 50 µm. Clinical chemistries of **(c)** Serum creatinine (CRE), **(d)** blood urea nitrogen (BUN), **(e)** calcium, and **(f)** alanine transaminase (ALT) levels in pigs pre-implantation and post-2-week implantation in the colon. Statistics are calculated by Paired T-test analysis (N=5). ns = not significant.

**Figure 10.**
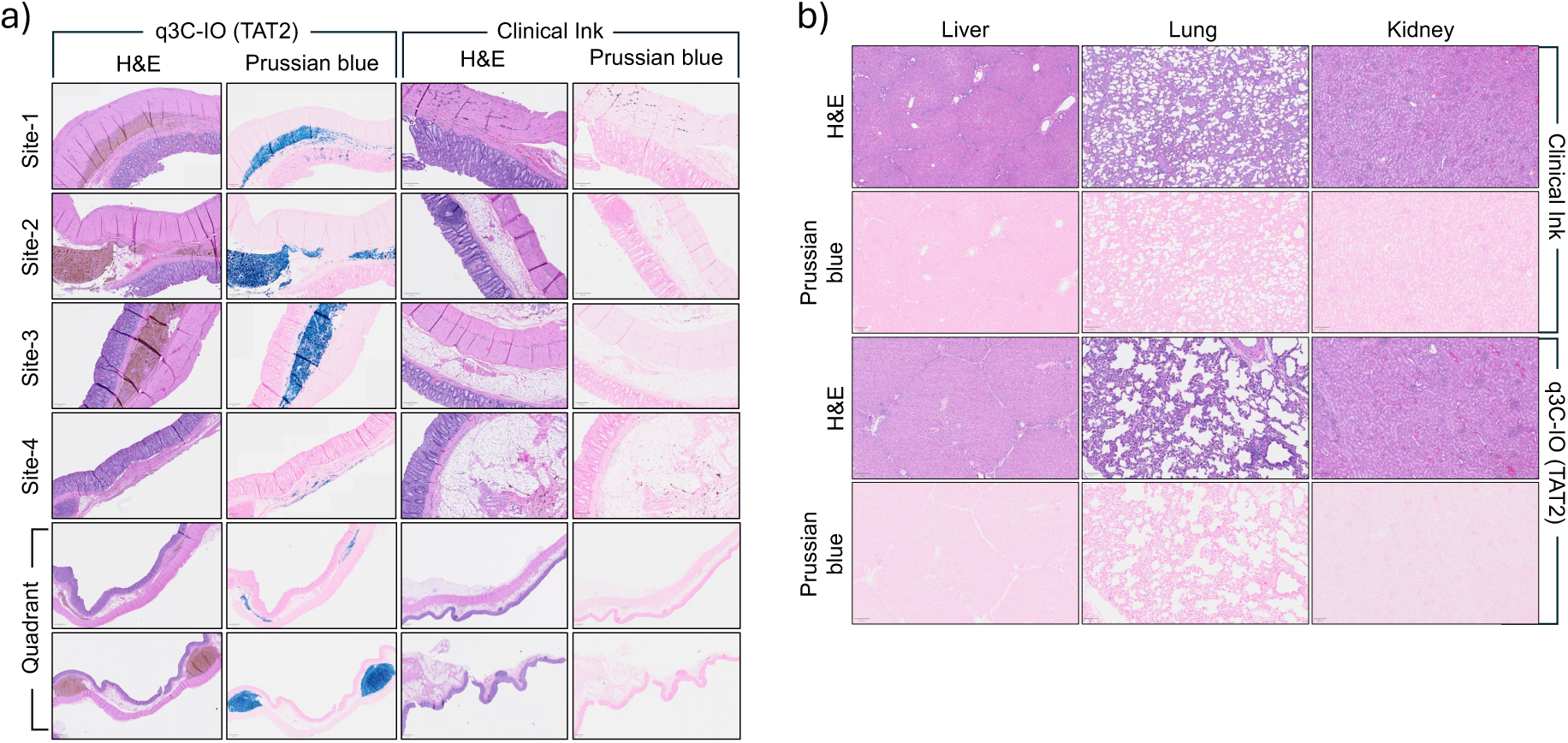
a) Representative histology images of H&E and Prussian blue staining of colon tissue administered with IO, hC-IO, q1C-IO & q3C-IO TAT2 ink and clinical inks after 65 days. (n=3) **b)** Representative H&E histology of pig end-organs 65 days post-implantation of clinical ink and TAT2 inks. No evidence of fibrosis or inflammation in the lung, inflammation or necrosis at the hepatic triad, or glomerulitis, capillaritis, or tubulitis in the kidney cortex was observed by blinded reading.

### Magnetic Resonance Imaging (MRI) using TAT2 Inks

Iron oxide (Fe_3_O_4_) nanoparticles are widely used for preclinical and clinical MR imaging and for the treatment of anemia (*70*). In MRI, it is crucial to evaluate relaxivities *r*_2_ and *r*_1_ (to shorten relaxation times T2 and T1, respectively), to improve the detection sensitivity of the MR contrast agent. Relaxivity quantifies how effectively a contrast agent enhances the relaxation rates of tissue water (R1=1/T1 or R2 =1/T2). Higher relaxivity means a more efficient contrast agent, which can induce a more significant change in relaxation rates (and consequently image intensities) at a given concentration (*71*).

The present study determined *r*_2_ and *r*_1_ values for IO and q3C-IO formulations under a 9.4 T magnetic field by assessing T2 and T1 relaxation times across varying Fe concentrations in each formulation (**Figure 11a**). Using a linear fit function, *r*_2_ and *r*_1_ relaxivities of q3C-IO were calculated as 242 ± 0.5 and 0.6 ± 0.05 mM^-1^ s^-1^, and *r_2_* and *r_1_* relaxivities of IO were calculated as 303 ± 0.5 and 0.7 ± 0.06 mM^-1^ s^-1^ (**Figure 11b**). The *r*_2_ values of both particles are higher than ferumoxide, ferucarbotran, and ferumoxtran, which are T2 contrast agents used in clinical studies. (*72, 73*) The relatively high relaxivities of q3C-IO and IO TAT2 inks can allow for better MR detection at lower concentrations. The *r*_1_ values of q3C-IO and dextran-coated IO are lower than clinically available T1 contrast agents, which is advantageous for use as a T2/T2* agent. Subcutaneous TAT2 inks (50 μL of 1 mg/mL IO and q3C-IO) implantation in a postmortem mouse in a 3T scanner demonstrated a strong T2 contrast signal (**Figure 11c**) that dominates even on a T1-weighted scan at clinical fields. The T2 contrast performance of TAT2 inks was similar to that of Ferumoxytol, a FDA-approved iron-based MRI contrast agent (*74*). However, no signal was observed from the clinical ink, a carbon-based suspension not expected to exhibit MR contrast.

**Figure 11.**
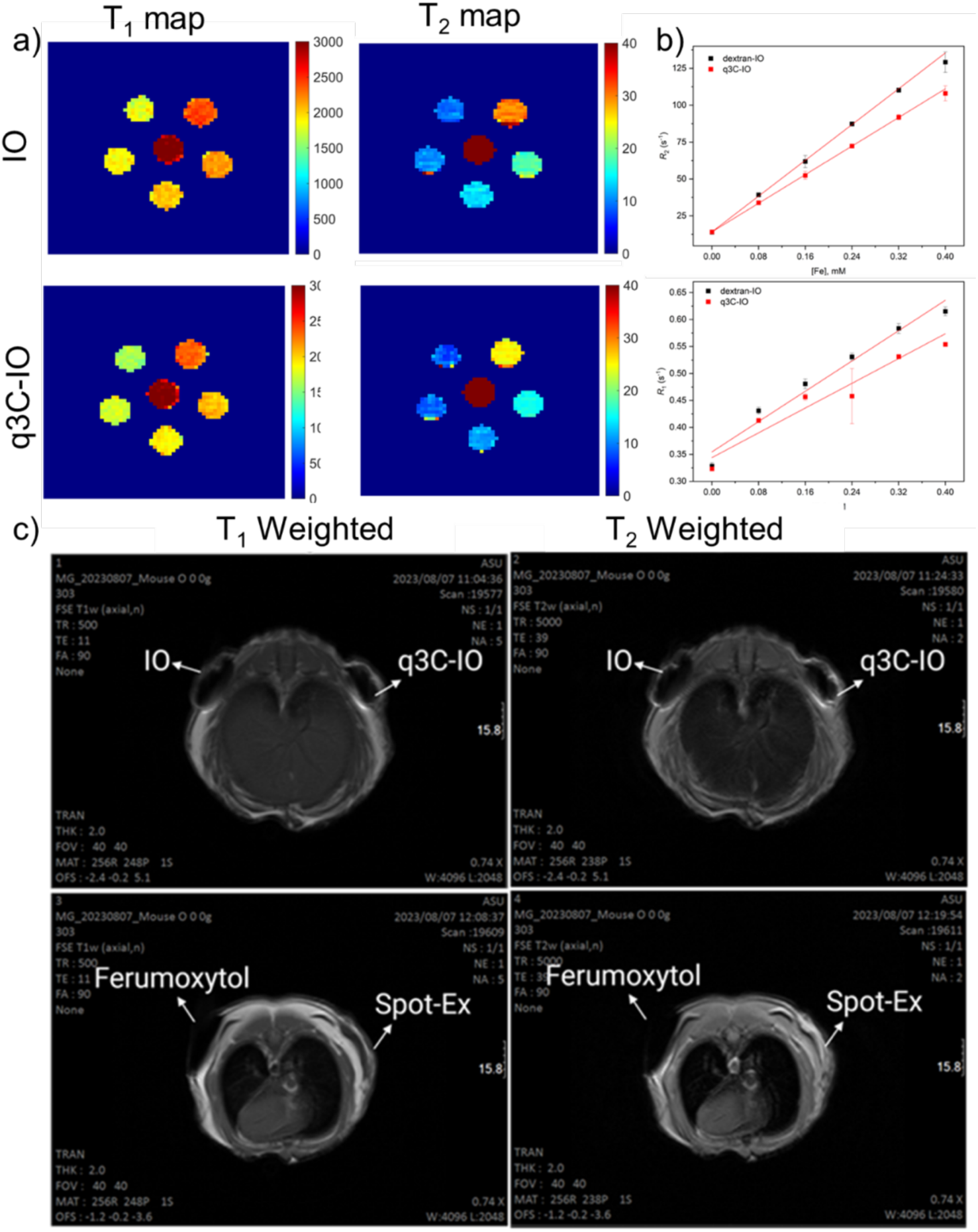
a) T_2_ and T_1_ maps of IO and q3C-IO in ms at concentrations of Fe (0, 0.08, 0.16, 0.24, 0.32, 0.4 mM) in 9.4T field. **b)** Relaxivity measurement of IO and q3C-IO. **c)** MRI images of postmortem mice injected with q3C-IO, IO, clinical ink, and Ferumoxytol in the subcutaneous region. Fast Spin Echo (FSE) images were captured using a 3T preclinical MRI scanner (representative of n=4 / group).

## Discussion

Preoperative endoscopic marking of colorectal lesions and polyps using the only FDA-approved carbon-based formulation has major limitations including extensive diffusion in the tissue resulting in poor spatial localization, and is associated with potential for peritoneal spillage, and lesional inflammation and fibrosis. We developed a biopolymer-coated iron oxide nanoparticle-based ink formulation as tissue-adhesive tattoos-generation 2 or TAT2 inks to overcome the several limitations of the current clinical ink. TAT2 inks demonstrated excellent dark contrast against white light and were stable for at least three months. TAT2 inks also demonstrated easy injectability through a standard endoscopic injection needle; the viscosity was similar to that of water and commercial ink. Cellular studies indicated no inflammatory activity of the TAT2 inks and studies with porcine intestinal tissues *ex vivo* indicated extensive diffusion of the commercial ink, whereas TAT2 inks maintained excellent spatial localization and contrast under white light. These characteristics were also noted when TAT2 inks were injected into the submucosa of live pigs, utilizing standard clinical endoscopic techniques. In contrast, extensive mucosal and serosal diffusion was noted with the clinical ink. TAT2 inks demonstrated excellent contrast under endoscopic light even for at least 65 days following injection. Histology and clinical chemistry demonstrated no adverse reactions or tissue toxicity with TAT2 inks, demonstrating their biocompatibility. TAT2 inks also demonstrated effective MR contrast properties, which enhances their utility particularly for multimodal noninvasive imaging and potentially expand the clinical scope of these inks beyond endoscopic visualization. Taken together, TAT2 inks demonstrate excellent biocompatibility, precise spatial localization, tissue contrast to endoscopic / white light, and MR contrast for multimodal imaging, making them effective next generation devices for endoscopic imaging. These inks also have the potential to be utilized in a broader scope such as for endoscopic ultrasound-guided marking of pancreatic tumors for precise surgical localization and for other disease states in Urology and Urogynecology.

## Materials and Methods

### Materials

HMC: High molecular weight chitosan (degree of deacetylation ≥ 75%; M_w_ = 310,000-375,000Da; purchased from Sigma), tHMC: high molecular weight chitosan ( purchased from Tidal vision), GTMAC: glycidyl trimethylammonium chloride (purchased from Sigma), dextran sulfate sodium salt (Mw = ∼40,000Da, purchased from Sigma), Iron(III)chloride (purchased from Sigma), Iron (II)chloride. Tetrahydrate (purchased from Sigma), Ammonia solution 25% (Suprapur^®^, purchased from Sigma), acetic acid (purchased from Sigma), Endotoxin free water (HyClone™ purchased from Cytiva). Spot^®^ Ex (Clinical ink, purchased from GI Supply – now Laborie, Camp Hill, Pennsylvania)

### Synthesis of quaternary ammonium conjugated chitosan derivatives (q1C, q2C & q3C)

HMC was conjugated to glycidyl trimethylammonium chloride (GTMAC) at three different degrees of conjugation: low (q1C), moderate (q2C) & high (q3C). Chitosan (HMC) 500 mg was dissolved in 40 mL of aqueous acetic acid solution (0.5% (v/v)) in a 100 mL round-bottom flask, and the pH was adjusted to 6.1-6.3, using an appropriate amount of 1N NaOH. The temperature was maintained at 60°C, placing the container in an oil bath for the duration of the reaction, which was carried out with shaking at 900-1000 rpm with a magnetic stirrer attached to a hot plate. To this vigorously stirring solution, three drop-wise additions of 87.5 μL (0.67 mmol), 265 μL (2.68 mmol), or 875 μL (10.7 mmol) of GTMAC were performed at 2.5 h time intervals in independent reactions to get q1C, q2C, and q3C, respectively. The stirring was continued at the same temperature with the same rpm for ∼24 h. Then, each solution was transferred to a 3.5 kDa dialysis membrane and purified against nanopure water for three days by changing the media 7-8 times. The retentate was lyophilized (LABCONCO^®^ Lyophilizer) to obtain the purified product in powder form.

### Synthesis of dextran-coated Iron oxide nanoparticles (IO) using the co-precipitation method

Dextran sulfate (40,000 g/mol; 1 g) dissolved in 60 mL endotoxin-free water was added to a 1L triple-necked round-bottom flask. A solution of FeCl_3_ (1.8 g, 11 mmol) and FeCl_2_.4H_2_O (1.26 g, 6.4 mmol), dissolved in 60 mL endotoxin-free water, was added dropwise to the dextran sulfate solution under magnetic stirring at 300 rpm, at room temperature under nitrogen gas. After ten minutes, 200 mL of NH_4_OH (7% v/v) was added dropwise for about 15 minutes and stirred for another 15 minutes at the same rpm and temperature. Then the temperature was raised to a range between 65°C-70°C, by placing the reaction vessel in an oil bath, and stirring was continued for about 30 minutes. All contents were cooled to room temperature, transferred into 50 mL centrifuge tubes, and centrifuged (6000g, 4°C, 20 min). The supernatant was discarded, and the nanoparticle precipitate was resuspended in endotoxin-free water. This procedure was repeated at least three times to ensure the complete removal of the NH_4_OH remaining with the nanoparticles. Finally, nanoparticles were resuspended in 30 mL of endotoxin-free water, and the pH of ∼9.5 was used to measure the concentration with the solvent evaporation method, in which 200 µL of nanoparticle liquid was placed in a microcentrifuge tube and evaporated the liquid in a vacuum centrifuge (Eppendorf^®^ Vacufuge Plus) at 40 °C for about 3-4 h.

### Preparation of chitosan-coated IO nanoparticles (C-IO, TAT2 ink)

tHMC solution (2%, w/v, pH=3.5), q1CH solution (2%, w/v, pH=3.5), and q3CH solution (2%, w/v, pH=3.5) were neutralized individually using 1N NaOH solution in order to adjust the pH to 6. Endotoxin-free water was then added to adjust the concentration of these biopolymers to 10 mg/mL (1%, w/v). The dispersion concentration containing dextran-coated iron oxide (IO) nanoparticles was also adjusted to 10 mg/mL using endotoxin-free water. Equal volumes (5 mL each) of IO and different chitosan solutions were mixed in separate sterile Falcon tubes and mixed under vortexing at 2500 rpm for ∼12 hrs at room temperature. The pH of the resulting black-colored suspension was adjusted to ∼6.5 using 1N NaOH solution. This C-IO dispersion was centrifuged (6000 *g*, 4°C, 20 min) to remove excess chitosan or chitosan-derived biopolymers. Upon discarding the supernatant, the pellet containing the black-colored ink particles were resuspended in endotoxin-free water and vortexed (for 5 minutes) to promote further mixing. This procedure was repeated three times to ensure complete removal of free (excess) chitosan (confirmed using the ninhydrin assay described below) from these dispersions. After the third round of centrifugation, the pellet containing C-IO particles was resuspended in 5 mL of endotoxin-free water to obtain a final TAT2 ink dispersion at 10 mg/mL concentration. Four different TAT2 inks, namely IO, hC-IO, q1C-IO, and q3C-IO, were generated and used in all the studies.

### ^1^H NMR Spectroscopy

^1^H NMR Spectra were recorded with a Varian Wide Bore Walkup spectrometer (400 MHz) at 353 K. All chitosan and chitosan derivative samples were dissolved at 2% v/v% DCl in D_2_O (10 mg of chitosan in 1 mL of DCl:D_2_O). Spectra were acquired for three independent batches of polymer samples.

### FT-IR Spectroscopy

Lyophilized powder of chitosan and chitosan derivatives were used for FT-IR spectroscopy. Spectra were acquired in transmittance mode using an FT-IR spectrometer (Perkin Elmer basic) equipped with a Smart iTR attenuated total reflectance (ATR) sampling accessory and diamond crystals. In case of TAT2 ink, the samples were dried, and the powder was used for FT-IR spectroscopy. Three independent samples of each ink were scanned at least three times, and the average of these spectra was used.

### Ninhydrin assay

Ninhydrin assay was carried out to quantify the number of primary amines in chitosan and its derivatives based on the manufacturer’s protocol (2% solution of Ninhydrin Reagent, SIGMA-N7285). 15 µL (1% w/v of chitosan solution) of each sample was mixed with 0.05% of aqueous acetic acid solution to make 200 µL of the mixture solution, which was then added with 100 µL of ninhydrin reagent. The resulting solution was placed in water bath at ∼100°C for 10 min. In this assay, the presence of reactive amines is indicated by a purple color of the solution. The absorbance of the final solution was measured at 570 nm. The amine content was determined by comparing the absorbance of the sample solution to a calibration curve generated using glycine standards.

### Degree of Quaternization (DQ%) of qC

The Mohr titration of chloride ions with a standard silver nitrate solution was used to determine the content of GTMAC functionalized d-glucosamine in chitosan. The conductometric titration was employed, with 0.05 N silver nitrate solution as the titrant and 25 mL of 0.2 wt% % synthesized qC serving as the analyte. After reaching the equivalence point between Ag^+^ and Cl^-^ions, the volume of AgNO3 was recorded. The degree of quaternization (DQ) is calculated using Equation 1:

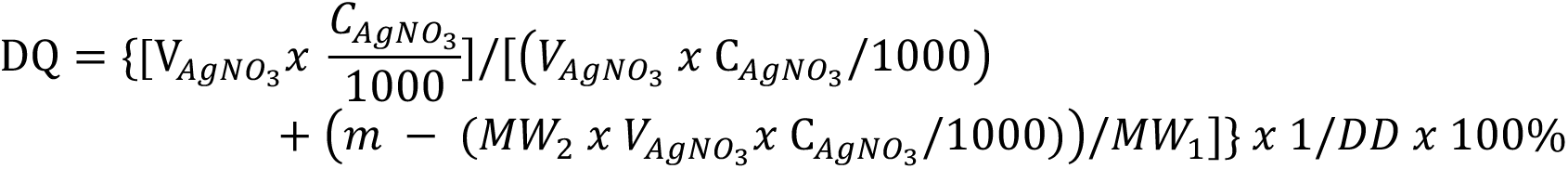

In equation 1, DD represents the degree of deacetylation, while MW_1_ and MW_2_ denote the molecular weights (g/mol) of d-glucosamine and N-(2-hydroxy) propyl-3-trimethyl ammonium chitosan chloride, respectively. V_AGNO3_ refers to the volume (mL) and 𝐶_AGNO3_ to the concentration (mol/L) of silver nitrate utilized in conductometric titration. The variable m indicates the mass (g) of qC dissolved in solution (*38*).

### Hydrodynamic diameter and zeta potential measurements

Hydrodynamic diameter and zeta potential values of candidate TAT2 inks were determined using a Zetasizer LAB (Malvern Instruments). Dispersions of TAT2 inks (100 μL of 10 mg/ml) were diluted to 5 mL stocks using Nanopure water (NPW) and analysed for hydrodynamic diameter using the side scatter method in triplicate. The average of these measurements was reported. One mL of the same stock ink dispersion was injected into a DS1070 zeta cuvette for zeta potential measurements, and an average of the three independent measurements was reported.

### Stability of TAT2 inks

TAT2 inks were stored at 4 °C in a degassed desiccator protected from light. Colloidal stability was tested by DLS verifying hydrodynamic diameter and zeta potential parameters (hydrodynamic diameter and zeta potential) every week for the indicated time.

### Scanning Electron Microscopy (SEM) and Electron Dispersion X-ray Spectroscopy (EDX)

Size and elemental analysis of TAT2 inks were analyzed using focused ion beam scanning electron microscopy (FIB-SEM; Auriga-Zeiss SEM). TAT2 inks were placed on double-sided carbon adhesive tape attached to an aluminium stub. Scanning electron microscopy was performed at a voltage of 5 kV, and EDS was carried out at 20 kV.

### Transmission Electron Microscopy (TEM)

TAT2 inks (8 μL) were added to a copper grid coated with formvar/carbon and dried at room temperature overnight. Imaging was performed on a Philips CM200 or JEOL TEM/STEM 2010F microscopes.

### ***X-*** ray diffraction (XRD)

XRD patterns were acquired using a Rigaku SmartLab high-resolution XRD spectrometer with a CuKα radiation source (A=1.54 A) at 40 kV and 30 mA scanning from 5° to 65° at a scan rate of 1° per minute.

### Viscosity measurements

Rheology test was performed using the flow sweep method by maintaining the torque force from 0.1 N to 50 N after placing 0.5 mL of each TAT2 ink, water, or Clinical ink upon a 20 mm sandblasted parallel plate connected to a rheometer (TA Instruments, Discovery HR30, New Castle, DE, USA) at 25 °C.

### Gel clot assay

ToxinSensor™ Gel Clot Endotoxin Assay Kit (GenScript) was used to determine the presence of endotoxin in TAT2 ink dispersions used in in vivo studies. The assay was performed as per the manufacturer’s protocol provided with the kit. All the TAT2 candidates were tested before any in vivo study within 24 h of the injection procedure. 100 µL of each TAT2 ink sample at different concentrations (10, 5, 2.5, 1.25, and 0.625 mg/mL) was taken into endotoxin-free glass vials provided with the kit, which were added with 100 µL of LAL (Limulus amoebocyte lysate) reagent which can form a clot after incubating the vials at 37 °C for one hour. If there is an endotoxin in any of the samples, the clot formation can be seen by flipping them up and down, which confirms that the sample contains ≥0.25 EU/mL of endotoxin. *E. coli* Endotoxin standard (5 EU/mL) is the positive control, and endotoxin-free water is the negative control in the assay.

### ICP-MS analysis of TAT2 inks

A 0.1 mL sample from 1 mg/mL stock of a TAT2 ink dispersion was weighed in a Teflon vial. Then, 0.25 mL of metal-free nitric acid and 0.75 mL of metal free hydrochloric acid were added to the ink dispersion. The vial was capped and placed on a hot plate overnight. The sample was then allowed to cool to room temperature before being transferred to a metal-free centrifuge tube. Nanopure water was added to the tube to bring the volume up to 10 mL, and the weight was taken again. Then, a 0.1 mL aliquot of this stock solution was taken and weighed in a metal-free centrifuge tube. 2% metal-free grade acid was added to a final volume of 10 mL, and the weight was retaken. This second solution is measured on the ICP-MS.

### In vitro Inflammation assay using J774-DUAL reporter macrophages

*In vitro* assessment of inflammatory transcription factor activity was performed using the J774-DUAL mouse macrophage reporter cell line (Invivogen, #j774d-nfis), a cell line derived from the parental J774A.1 line that preserves critical innate immune and inflammatory responses(*75–78*). J774-DUAL cells are engineered to stably contain coding sequences for secreted Lucia luciferase under the control of an IRF3 transcription factor binding site (minimal ISG54 promoter with five IFN-stimulated response elements) and for secreated alkaline phosphatase (SEAP) under the control of a NFΚB transcription factor binding site (minimal IFN-β promoter, five copies of the NFΚB consensus binding site, and three copies of the c-Rel binding site). Cells were maintained in DMEM containing 10% heat-inactivated FBS (Seradigm #1500-050H) and 1% penicillin-streptomycin (ATCC #30-2300), with further supplementation by selective antibiotics Blasticidin (5 µg/mL) and Zeocin® (100 µg/mL). Selective antibiotics were omitted when seeding cells for experiments. Cells were fed twice weekly and passaged by scraping.

For the reporter assay cells were passaged and viability was evaluated using the 0.4% Trypan Blue dye exclusion assay on an EVE™ Plus Automated Cell Counter (NanoEnTek) before resuspension at a density of 2.8×10^5^ cells/mL in medium. Cells were only used if viability was at least 95%. Prior to cell seeding, 20 µL of a 10X concentration of material, TAT2 ink, positive control (1 µg/mL E. coli O111:B4 LPS), or medium were pre-dispensed in each well of a tissue culture-treated 96-well plate. Resuspended cells were dispensed in each well in a volume of 180 µL (∼50,000 cells/well final), diluting the dispensed condition to 1X working concentration. Cells were incubated at 37°C/5% CO_2_ for 24 hours. After placing cells in the incubator, same-day endotoxin levels were evaluated using the ToxinSensor™ gel clot assay (Genscript # L00351), a limulus amoebocyte lysate (LAL) assay and confirmed to be endotoxin-free at <0.25 EU/mL. After 24 hours incubation 20 µL of conditioned supernatant was transferred to a clear-wall 96-well plate with 180 µL of 1X working concentration of QUANTI-Blue SEAP assay reagent, incubated at 37 °C for 30 minutes, and OD630 was measured on a Biotek Synergy H2 multimode plate reader as a readout of NFΚB activity. Separately, 30 µL of conditioned supernatant was transferred to a opaque, white-walled 96-well plate and 50 µL of QUANTI-Luc Lucia assay reagent was added to each well using a multichannel pipette with immediate luminescence reading at 0.1 seconds integration on a Biotek Synergy H2 multimode plate reader as readout of IRF3 activity.

### Ex vivo studies on porcine colon tissue using TAT2 inks

Fresh porcine large intestine was purchased from Animal Technologies. The intestine was cleaned with DI water 2-3 times and cut into uniform rectangular pieces (5 cm x 3 cm), which were stored in saline under 4°C until further use. Then a portion of the intestinal tissue was placed on an inverted weighing boat and ∼50 µL of a candidate TAT2 ink or the clinical ink was injected (as shown in **Figure 4a**) into the submucosal layer with an insulin syringe (30G) at ∼10-degree angle with tissue surface. The tattoo/mark was visualized immediately or at a specific time point after storing in saline at room temperature. Images of the ink in tissue were acquired using a cell phone camera placed within an enclosed, house-made black box imaging system, which was assembled to avoid any lighting artifacts and maintain uniform light conditions. All images were calibrated and analyzed using a custom macro in FIJI/ImageJ to measure contrast and area of each spot created with the TAT2 inks or clinical ink. Briefly, the macro used in-image scaling (via a ruler) to apply calibration to the image, processed thresholds to identify spot on the basis of brightness/darkness, and performed grey scale measurements of mean intensity on both the spot region as well as background regions. All images were processed in a linear manner and investigator feedback was only prompted for identification of the gross spot region and the gross background regions with no investigator control of thresholding.

### In vivo biocompatibility of TAT2 ink in mice

Balb/c mice were purchased from Jackson Laboratories, ME, USA. At least one animal from the opposite sex was used in each group. All procedures were approved by the ASU Institutional Animal Care and Use Committee (IACUC). Mice, 13 weeks of age, were anesthetized by an intraperitoneal (i.p.) injection (100 µL) of a cocktail containing ketamine (120 mg/kg) and xylazine (6 mg/kg). The dorsum was shaved using electric clippers, and the surgical area was sterilized three times by cyclic application of 2% chlorhexidine gluconate and 70% ethanol. Two 50 µL injections of different TAT2 inks or clinical ink were administered subcutaneously, ∼1 cm apart, to each animal as shown in **Fig. 5A** and the images were collected by phone digital camera and mice were returned to their cages. Mice were monitored for up to 4 weeks with regular photographic documentation of injection sites. At 4 weeks, the mice were euthanized, the dorsal skin was removed, and the subcutaneous region was imaged. Tissues were collected in formalin and processed into FFPE for histopathology by Hematoxylin and eosin (H&E) staining. Slides were imaged on an Olympus BX43 microscope fitted with a DP74 camera operated by cellSens Dimensions software.

### Endoscopic submucosal colonic injections in live pigs – 2 week study

Four Yorkshire pigs, two male and two female (age: 3 – 4 months old, 50 - 60 kg), were purchased from Premier BioSource, California, USA. All procedures were approved by the Institutional Animal Care and Use Committee (IACUC) at the Mayo Clinic. Pigs were acclimatized for at least a week before the procedure, administered a fleet enema twice on each consecutive day, two days prior to the procedure, and were subsequently kept on a liquid diet. Two hours before the procedure, animals were anesthetized by intramuscular (i.m.) injection of telazol (5 mg/kg), xylazine (2 mg/ kg), and glycopyrrolate (0.02 mg/kg). Colonoscopy was performed through rectum using a therapeutic gastroscope (TJF 190, Olympus, America, Center Valley, PA) and ∼1 mL of each TAT2 ink or clinical ink was injected approximately 5 cm apart from each other. Each pig, therefore, received a total of 5 different tattoos. Pigs were maintained under isoflurane for the duration of the colonoscopy procedure. Following endoscopy, videos and images were recorded, and the pigs were returned to their housing facility and observed for 2 weeks. On day 12 and day 13 two fleet enemas were administered, and the pigs were maintained on a liquid diet. On day 14, two hours before the procedure, animals were anesthetized by intramuscular (i.m.) injection of telazol (5 mg/kg), xylazine (2 mg/kg), and glycopyrrolate (0.02 mg/kg). Colonoscopy was again carried out as described previously, and images and videos were recorded. Pigs were euthanized by administering (DRUG name), and tissue specimens were acquired and stored in formalin for subsequent histological analyses.

### Endoscopic submucosal colonic injections in live pigs – 65-day study

Four Yorkshire pigs (age: 2 months old, 20 kg; all male) were evaluate with clinical ink and three Yorkshire pigs (age: 2 months, 20 kg; 2 male and 1 female) for qC-IO evaluation were purchased from Premier BioSource, California, USA. All procedures were approved by the Institutional Animal Care and Use Committee (IACUC) at the Mayo Clinic. Pigs were acclimatized for at least a week before the procedure, administered a fleet enema twice on each consecutive day, two days prior to the procedure, and were subsequently kept on a liquid diet. Two hours before the procedure, animals were anesthetized by intramuscular (i.m.) injection of telazol (5 mg/kg), xylazine (2 mg/ kg), and glycopyrrolate (0.02 mg/kg). Colonoscopy was performed through rectum using a therapeutic gastroscope (TJF 190, Olympus, America, Center Valley, PA) and ∼1 mL of each TAT2 ink or clinical ink was injected approximately 5 cm apart from each other. Each pig, therefore, received a total of 5 different tattoos. Pigs were maintained under isoflurane for the duration of the colonoscopy procedure. Following endoscopy, videos and images were recorded, and the pigs were returned to their housing facility and observed for 65 days. On day 38 and day 39 two fleet enemas were administered, and the pigs were maintained on a liquid diet. On day 40, two hours before the procedure, animals were anesthetized by intramuscular (i.m.) injection of telazol (5 mg/kg), xylazine (2 mg/kg), and glycopyrrolate (0.02 mg/kg). Colonoscopy was again carried out as described previously, and images and videos were recorded. Pigs were returned to their housing and monitored for another 25 days. On day 63 and 64, pigs were administered fleet enemas, anesthetized as above on day 65, and a colonoscopy was performed. Pigs were euthanized by administering (DRUG name), and tissue specimens were acquired and stored in formalin for subsequent histological analyses.

### Histology

At follow-up, mouse skin and pig colon, liver, kidney, and lung were collected in 10% neutral-buffered formalin and fixed at room temperature for 3-7 days. Tissues were processed manually by washing with two PBS exchanges, dehydrating through a graded alcohol series (40%, 70% x2, 90% x2, 100% x2), clearing with three changes of 100% xylene, partially perfusing in 1:1 xylene:paraffin, and finally perfusing through three exchanges with 100% paraffin as described previously(*79, 80*). Steps involving paraffin were performed at 60 °C, all other steps were performed at room temperature. Following paraffin perfusion, tissues were embedded in Paraplast Plus (EMS Diasum #19217). Single 5-µm sections were cut with an Accu-Cut® SRM™ 200 Rotary Microtome (Sakura Finetek USA) and captured on charged glass slides (Hareta) in a floating water bath maintained at 42.4 °C (IHC World #XH-1003). Slides were dried overnight at 37 °C and stored at room temperature until staining.

For staining, slides were pre-incubated at 60 °C for 30-60 minutes to soften paraffin. Slides were deparaffinized through two exchanges of 100% xylene and rehydrated through graded alcohol (90%, 70%) into tap water. For H&E staining, slides were submerged in Gill No. 2 Hematoxylin (Sigma Aldrich #GHS232) for 3 minutes, differentiated by 12 quick dips in acid alcohol (0.3% hydrochloride acid in 70% ethanol), and blued in a solution of ammonium water (0.2% ammonium hydroxide in distilled water) for 30 seconds. Slides were then submerged in 0.5% Eosin Y (Sigma Aldrich #318906) prepared in distilled water acidified with 0.2% glacial acetic acid for 4 minutes followed by combined washing/dehydration in 90% and 100% ethanol exchanges. Slides were further dehydrated through two exchanges of 100% xylene, air-dried, and mounted with CytoSeal XYL (Richard-Allan/Thermo Fisher) and #1.5 cover glass. For Prussian Blue staining, slides were deparaffinized and rehydrated into tap water following the same procedure as for H&E. Rehydrated slides were submerged in a Prussian Blue working solution (equal parts 2% hydrochloric acid and 2% potassium ferrocyanide [EMS Diasum #26613-01] prepared just before use) in a coplin jar until brilliant blue staining was observed on the slides (approximately 1-2 minutes) followed by thorough washing in three exchanges of distilled water and dehydration and mounting as described for H&E. Samples were imaged on an Olympus BX43 upright microscope configured with an Olympus DP74 CMOS camera and operated by cellSens Standard Software (Olympus Corporation).

### Clinical Blood chemistry

Anticoagulated blood was collected pre-and post-procedure for all animals at the timem of initial ink placement and at terminal follow-up. Clinical blood chemistry was quantified on a Heska/Fuji Dri-Chem FDC4000V v1.2-PO1 and analyzed in Graphpad Prism by paired T-test.

### Relaxivity measurements using Phantom based MRI scanning

Relaxivities of q3C-IO TAT2 ink and IO nanoparticles were determined using a variable-field MR Solutions preclinical MRI scanner operating at 9.4 T. The relaxivity of the materials were studied using a home-made 3D printed phantom. This phantom has cylindrical geometry with a 30 mm diameter base and 19.2 mm height. A cavity is bored out of the main body to allow water flow and heat exchange with flowing water through inlets and outlets connected to a circulating water bath set to 37 °C to simulate physiological conditions. SIX holes are created at the top of the phantom to allow ∼1ml Eppendorf tubes to be placed in direct contact with the circulating water. These tubes contained six different concentrations of TAT2 inks and clinical ink in water at 0, 0.2, 0.4, 0.6, 0.8, 1 mg/mL, respectively. The concentration of iron in stock solutions was determined using ICP-MS and these concentrations were used to generate relaxivity plots. A multi-echo MRI sequence was used to measure T2 values of a coronal slice through the phantom using the parameters: slice thickness= 1 mm, TR=4000 ms, 60 echos with minimum TE=11 ms and echo spacing of 11 ms. MR images were analysed, which resulted in determinations of T2 values for each pixel. Map of the relaxation time T2 was computed on a pixel-by-pixel basis from least-squares fitting of the exponentially decaying signals (as a function of echo time) using a custom MATLAB code. Regions of interest (ROIs) were drawn on the maps around each vial to obtain the mean T2 for each and these were converted to relaxation rate R2 (=1/T2). The r_2_ relaxivity value was calculated as the slope of a linear fit of R2 vs Fe concentration. A Inversion Recovery Fast Low-Angle SHot (IRFLASH) sequence was used to measure T1 values of a coronal slice through the phantom using the parameters: slice thickness=1 mm, TR=10 ms, TE=4 ms, TI=50-2410 ms with intervals of 40 ms. Regions of interest (ROIs) were drawn on the maps around each vial to obtain the mean T1 for each and these were converted to relaxation rate R1 (=1/T1). The r_1_ relaxivity value was calculated as the slope of a linear fit of R1 vs Fe concentration.

### MRI performance of TAT2 in mouse cadaver subcutaneous implantation

Balb/C mice (6-8 weeks old) were euthanised using the protocol approved by Institutional Animal Care and Use Committee (IACUC) at ASU, and the dorsum was shaved using electric clippers. Two 50 µL injections of lead TAT2 ink (q3C-IO), IO, Ferumoxytol, or clinical ink were administered sub-cutaneously, ∼1 cm apart, to each animal as shown in **Fig. 5A** and the mouse was transferred to a preclinical Bruker MRI machine and MRS preclinical MRI machine operating at 3 Tesla (T).

## Acknowledgments

We acknowledge the use of facilities within the Eyring Materials Center at Arizona State University supported in part by NNCI-ECCS-1542160. We also gratefully acknowledge support from the Center for Procedural Innovation, Mayo Clinic in Arizona for the conduct of the porcine experiments.

## Funding

Flinn Foundation Translational Seed Grant (KR)

Mayo-Arizona State University Seed Grant (KR and RP)

## Author contributions

Conceptualization: RP and KR

Methodology: MG, JRY, SD, HS, MD, VDK, RP, KR

Investigation: MG, JRY, SD, HS, CW, VS, MD, TC Visualization: MG, JRY, VS, VDK

Supervision: VDK, RP, KR Writing—original draft: MG, JRY, KR

Writing—review & editing: MG, JRY, SD, HS, CW, VS, MD, TC, VKD, RP, KR

## Competing interests

KR is affiliated with Synergyan, LLC and Endotat Biotechnologies, LLC. JRY is affiliated with Vivo Biotechnologies, LLC and Endotat Biotechnologies, LLC. MG is affiliate with Endotat Biotechnologies, LLC. RP is affiliate with Endotat Biotechnologies, LLC.

## Data and materials availability

All data are available in the main text or the supplementary materials.

