## Supplemental Materials for "Tissue-Adhesive Endoscopic Tattoo Inks for Precise Marking and Surveillance of Gastrointestinal Lesions"

**This PDF file includes:**

Figs. S1 to S9
Movies S1 to S2
Table S1

**Other Supplementary Materials for this manuscript include the following:**

Movies S1 to S2

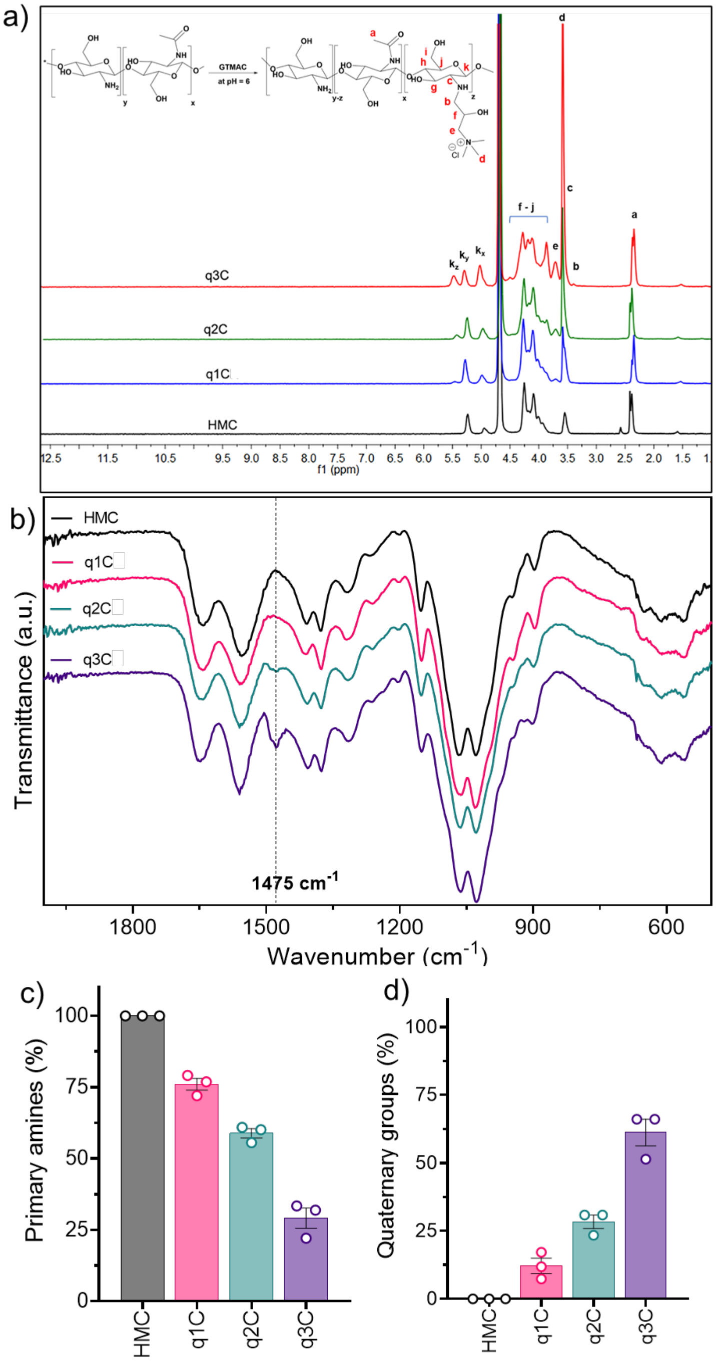


**Fig. S1. Synthesis and characterization of Biopolymers.** High molecular weight chitosan (HMC) and quaternized chitosan (q1C, q2C, and q3C) **a)** $^{1}$H-NMR spectra, **b)** FT-IR spectra of all the polymers used for TAT2 synthesis, with the peak at 1475 cm-1 highlighted for each polymer (N = 3 independent batches of polymers). c) Quantification of primary amines using the ninhydrin assay (N = 3). d) Quantifying trimethylammonium chloride (quaternary amine) groups in each polymer determined using conductometric chloride ion estimation using standard silver nitrate solution (N = 3).

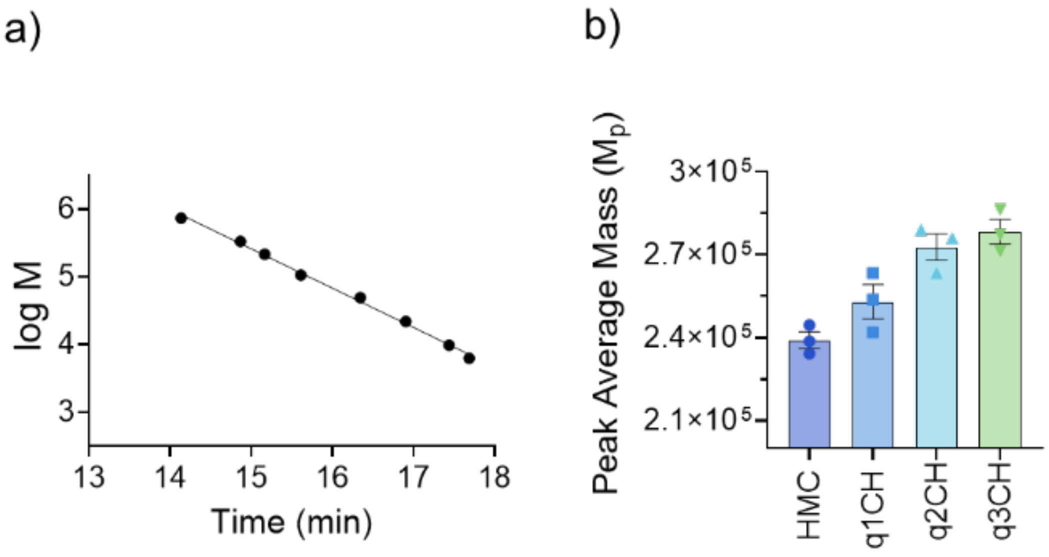


**Fig. S2: Gel permeation chromatography (GPC).** a) Molar mass calibration curve of pullulan standards on aquagel-OH MIXED-H 8 µm, 300 x 7.5 mm columns attached in series. Experimental analytical conditions: 0.5 M $NaNO_3$ + 0.01 M $NaH_2PO_4$ at pH 2 as mobile phase at 1 mL/min, RI detection, and room temperature (25 °C). An Agilent HPLC equipped with a 1200 Series refractometric detector was used for the experiments. The calibration equation is (log M) = -0.5809(time) + 14.128 ($R^2$ = 0.9967). b) Quantification of Peak average molecular weights (Mp) of chitosan samples HMC, q1CH, q2CH, and q3CH (N=3).

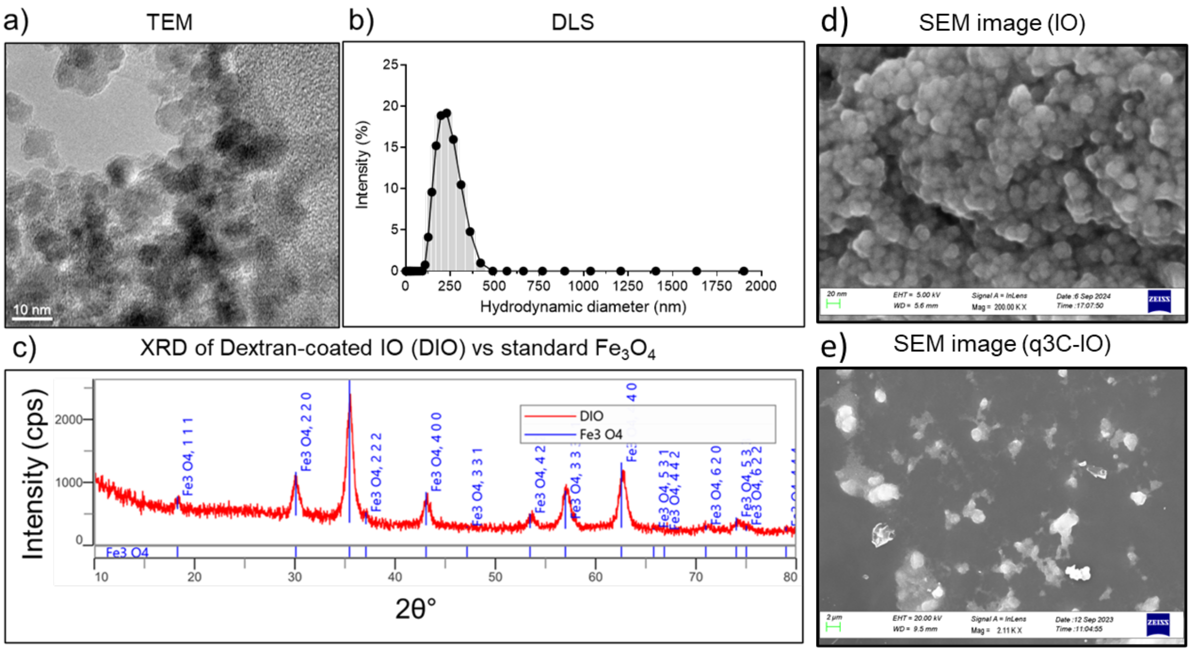


**Fig. S3:** a) Representative Transmission Electron Microscopic (TEM) image of IO (Scale bar 10 nm), b) Distribution of Hydrodynamic diameter of IO, c) X-Ray Diffraction (XRD) spectra of IO (DIO: Dextran-coated Iron Oxide – Red line spectra) nanoparticles in comparison with standard $Fe_3O_4$ (Blue line) crystallographic pattern. **d)** Scanning electron microscopy (SEM) images of IO (scale bar, 20 nm) and **e**) q3C-IO (scale bar, 2 μm) (bottom).

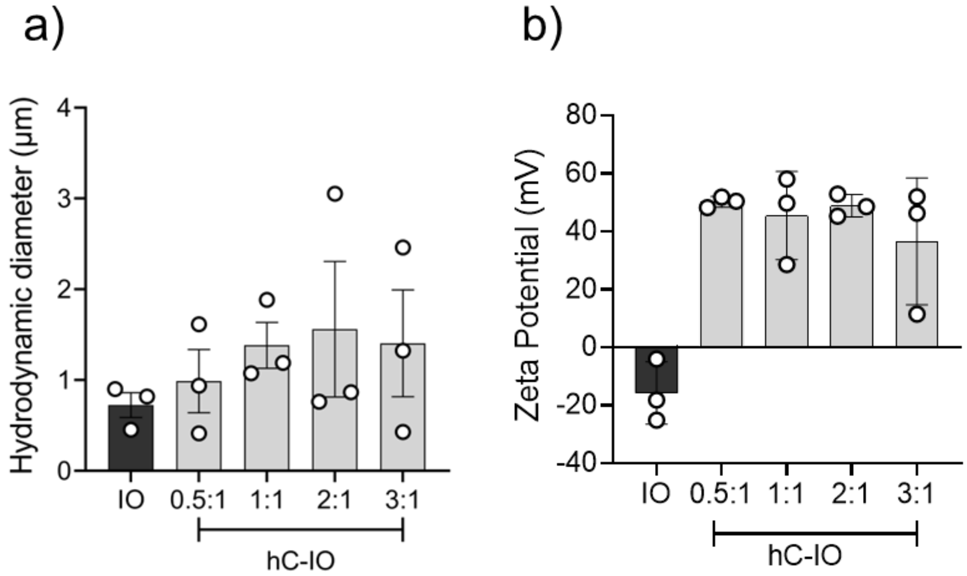


**Fig. S4: DLS Characterization of chitosan and IO ratio optimization:** a) Z-average hydrodynamic diameter and b) Zeta potentials of IO (pH = 8) and hC-IO (pH = 6.5) at different weight ratios of chitosan vs IO (hC-IO) (0.5:1, 1:1, 2;1, & 3:1).

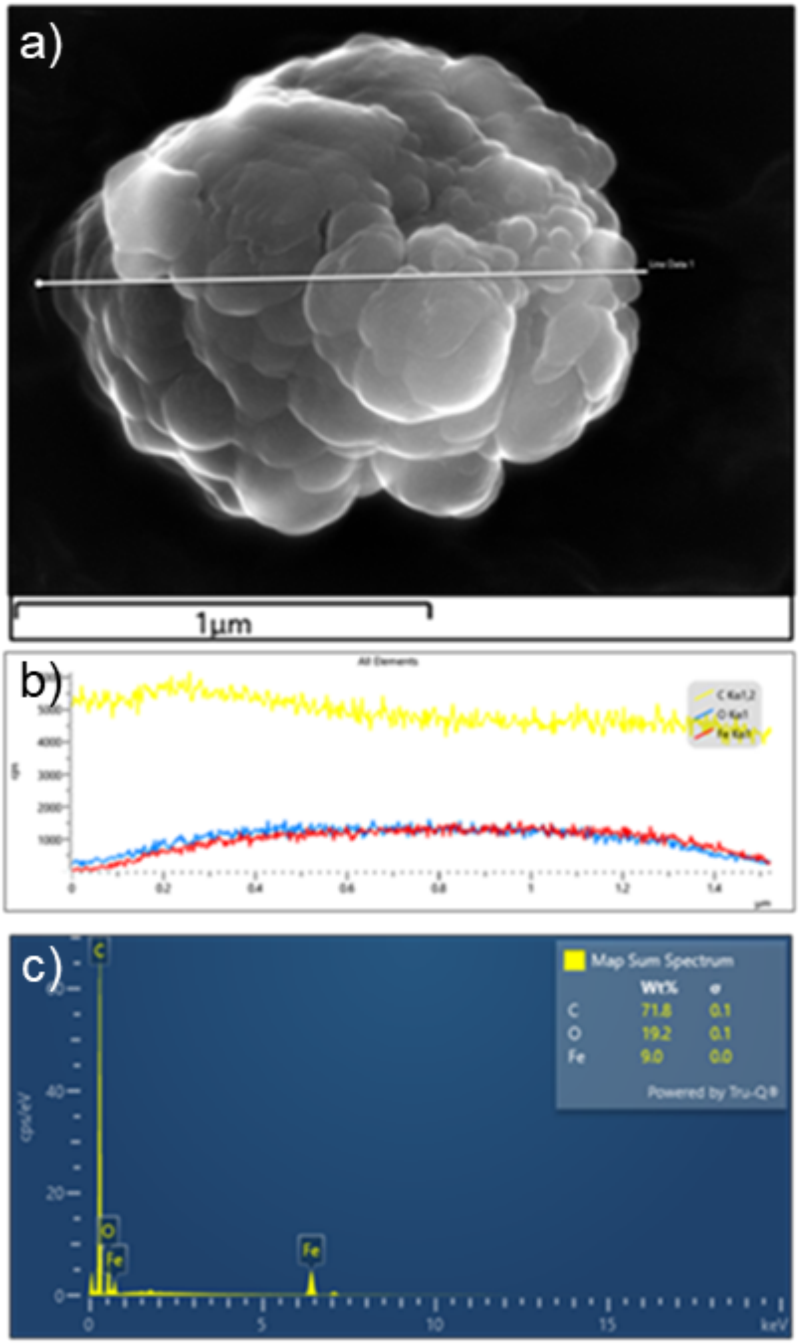


**Fig. S5:** a) SEM image of a single q3C-IO particle. b) Element-based spectra (carbon-yellow, oxygen-blue, and red-iron) generated by the line scan method in SEM-EDX, c) Energy dispersive X-ray method spectrum showing the weight ratios of carbon, oxygen, and iron present in a single particle of q3C-IO.

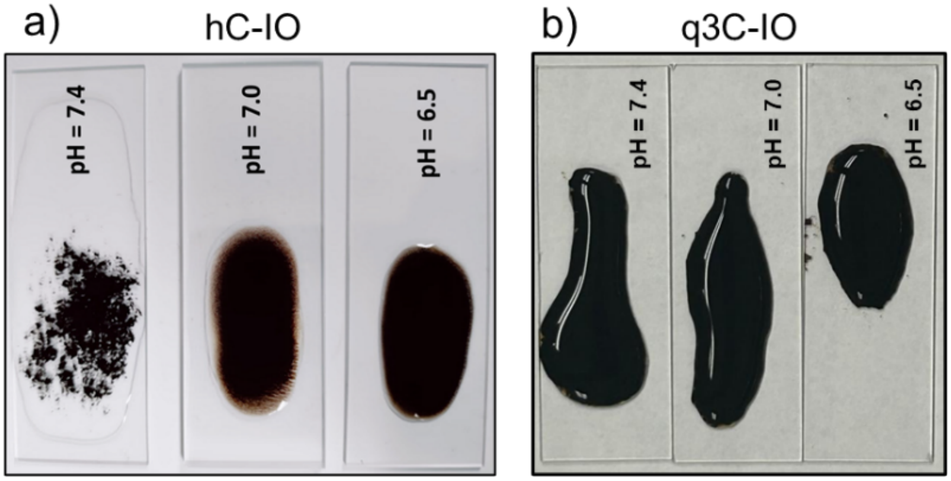


**Fig. S6**: Effect of pH on TAT2 dispersion: (a) hC-IO and (b) q3C-IO, each placed on a 2 × 4 cm glass slide using 0.5 mL per sample, synthesized at three different pH values, 6.5, 7.0, and 7.4.

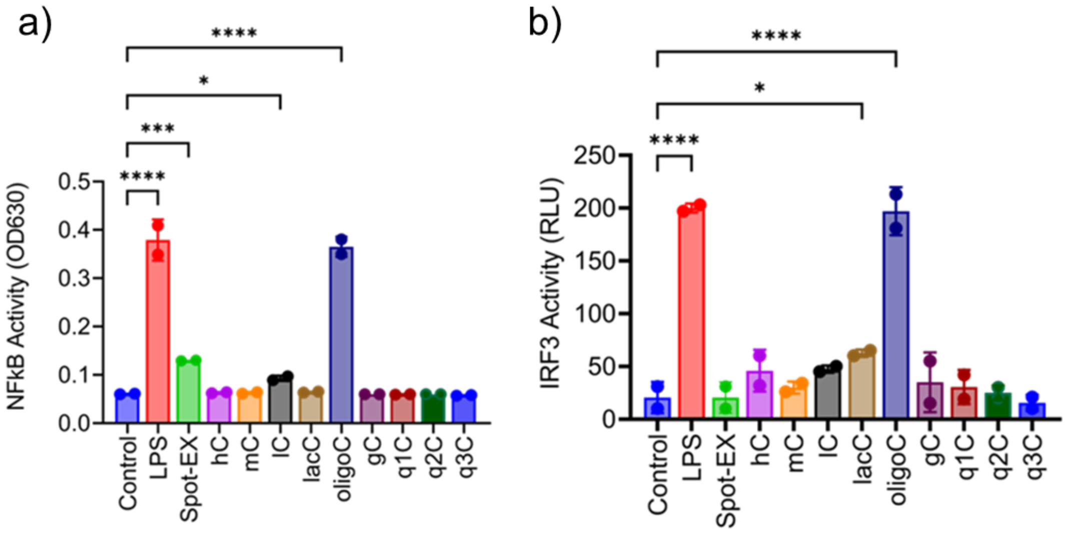


**Fig. S7**: (a) IRF3 activity measured by secreted luciferase and (b) NF-kB activity measured by secreted alkaline phosphatase of J774-DUAL$^{TM}$ mouse macrophage cell line treated with individual chitosan solutions after 24 hours of incubation. Lipopolysaccharide (LPS) was used as a positive control, and unstimulated cells as a negative control. Statistics were calculated as one-way ANOVA with Fisher's LSD using the untreated controls as the comparison group. *p<0.05, **p<0.01, ***p<0.001, ****p<0.0001.

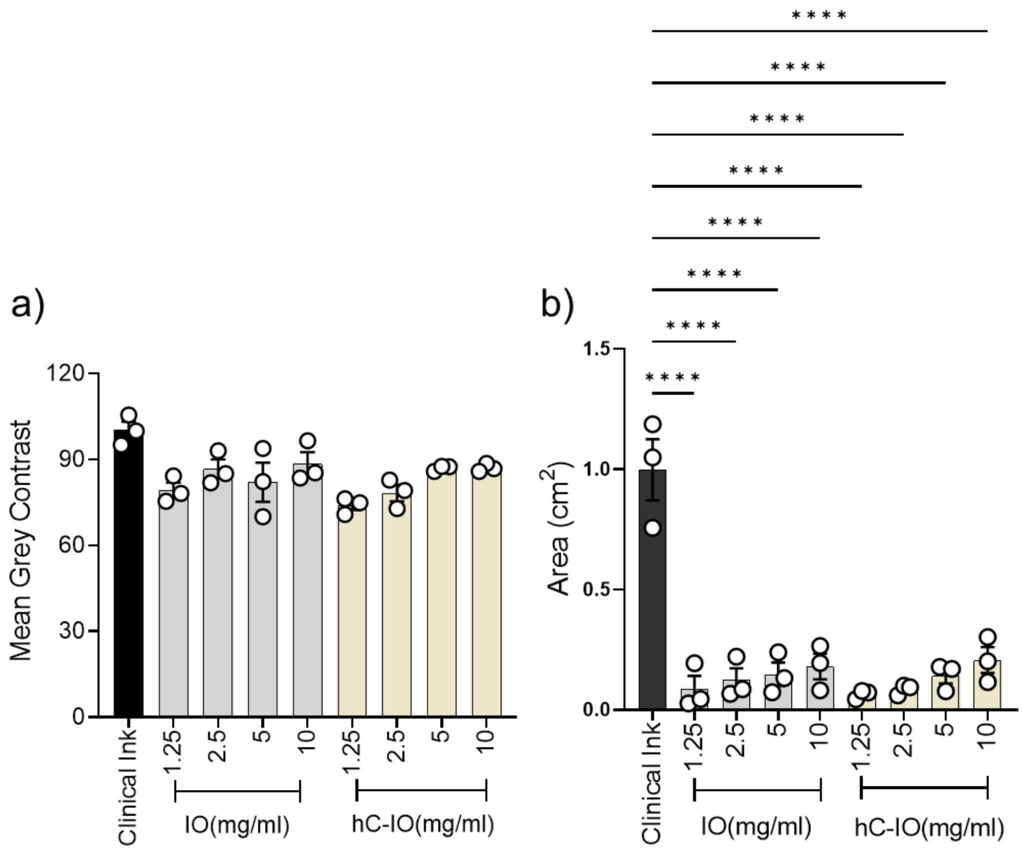


**Fig. S8**: Quantified ex vivo performance of TAT2 (IO and hC-IO) at different concentrations (1.25, 2.5, 5, & 10 mg/mL) of the particles by maintaining the same volume (50 µL) in terms of mean grey value of contrast (A) and Area ($cm^2$) of diffusion (B) in comparison to commercial formulation clinical ink (N=3). $*p<0.05$, $**p<0.01$, $***p<0.001$, $****p<0.0001$.

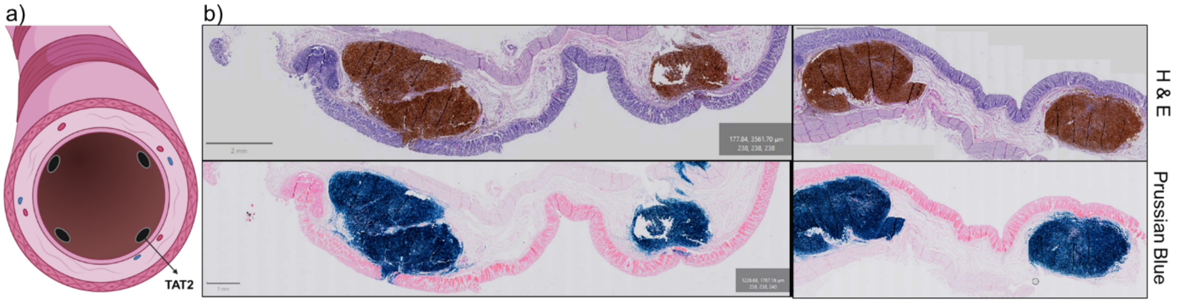


**Fig. S9**: a) Schematic shows the quadrant injection pattern in the colon, b) Histology images of all four locations of inks implanted with q3C-IO after 14-day residence in the colon and stained using H&E and prussian blue.

**Table S1.** Clinical chemistries after 65 days implantation with clinical ink or TAT2

| | ***BUN*** | | | ***CRE*** | | | ***Sr $Ca^{2+}$*** | | | ***ALT*** | | |
|---|---|---|---|---|---|---|---|---|---|---|---|---|
| **Day:** | **0** | **40** | **65** | **0** | **40** | **65** | **0** | **40** | **65** | **0** | **40** | **65** |
| *q3C-IO 1* | 6.4 | 11.9 | 14.2 | 0.8 | 1 | 1.2 | 7.9 | 9.6 | 10.5 | 42 | 49 | 46 |
| *q3C-IO 2* | 5.6 | 5.5 | 7.3 | 0.8 | 1.1 | 1.5 | 9.4 | 9.4 | 9.6 | 57 | 52 | 39 |
| *q3C-IO 3* | 5.3 | 4.9 | 5.9 | 0.8 | 1.1 | 1.5 | 8 | 8.2 | 8.9 | 33 | 33 | 31 |
| *Clinical 1* | < 5.0 | < 5.0 | 7.2 | 0.9 | 1 | 1 | * | 7.8 | 9.1 | 64 | 54 | 39 |
| *Clinical 2* | < 5.0 | < 5.0 | * | 1 | 1.3 | < 0.2 | * | 8 | * | 58 | 51 | 39 |
| *Clinical 3* | < 5.0 | < 5.0 | 7.1 | 0.9 | 1.1 | 1.3 | * | 8.8 | 9.4 | 53 | 40 | 30 |
| *Clinical 4* | < 5.0 | < 5.0 | 11 | 1 | 1 | 1.2 | * | 9.9 | 8.5 | 58 | 50 | 37 |

* value unavailable

**Movie S1.** Endoscopic Video of porcine colon on day 14 injected with clinical ink, IO, hC-IO, q1C-IO, and q3C-IO using each 1 ml for submucosal injection (N=5) followed by a quadrant injection using q3C-IO with 0.5 ml per injection (N=3).

**Movie S2.** Endoscopic Video of porcine colon on day 65 injected with q3C-IO using 1 ml at four different sites of porcine colon separated with 5 cm distance followed by a quadrant injection using 0.5 ml per injection (N=3).
